# ST6GAL1 sialyltransferase promotes acinar to ductal metaplasia and pancreatic cancer progression

**DOI:** 10.1101/2022.04.28.489561

**Authors:** Asmi Chakraborty, Nikita Bhalerao, Michael P. Marciel, Jihye Hwang, Colleen M. Britain, Isam E. Eltoum, Robert B. Jones, Katie L. Alexander, Lesley E. Smythies, Phillip D. Smith, David K. Crossman, Michael R. Crowley, Boyoung Shin, Laurie E. Harrington, Zhaoqi Yan, Maigen M. Bethea, Chad S. Hunter, Christopher A. Klug, Donald J. Buchsbaum, Susan L. Bellis

## Abstract

The role of aberrant glycosylation in pancreatic ductal adenocarcinoma (PDAC) remains an under-investigated area of research. In this study, we determined that the ST6GAL1 sialyltransferase, which adds α2,6-linked sialic acids to *N*-glycosylated proteins, is upregulated in patients with early-stage PDAC, and further increased in advanced disease. A tumor-promoting function for ST6GAL1 was elucidated using tumor xenograft models with human PDAC cells. Additionally, we developed a genetically-engineered mouse (GEM) with transgenic expression of ST6GAL1 in the pancreas, and found that mice with dual expression of ST6GAL1 and oncogenic KRAS^G12D^ have greatly accelerated PDAC progression and mortality compared with mice expressing KRAS^G12D^ alone. As ST6GAL1 imparts progenitor-like characteristics, we interrogated ST6GAL1’s role in acinar to ductal metaplasia (ADM), a process that fosters neoplasia by reprogramming acinar cells into ductal, progenitor-like cells. We confirmed that ST6GAL1 promotes ADM using multiple models including the 266-6 cell line, GEM-derived organoids and tissues, and an *in vivo* model of inflammation-induced ADM. EGFR is a key driver of ADM and is known to be activated by ST6GAL1-mediated sialylation. Importantly, EGFR activation was dramatically increased in acinar cells and organoids from mice with transgenic ST6GAL1 expression. These collective results highlight a novel glycosylation-dependent mechanism involved in early stages of pancreatic neoplasia.

## Introduction

Pancreatic ductal adenocarcinoma (PDAC) is an aggressive cancer with a dismal 5-year survival rate of less than 10% (1). Accordingly, there is a pressing need to elucidate the molecular mechanisms that underlie PDAC development. One of the earliest events contributing to pancreatic neoplasia is acinar to ductal metaplasia (ADM), a process in which acinar cells acquire a more ductal, progenitor-like phenotype (2-4). The disease then progresses through varying stages of premalignant lesions known as Pancreatic Intraepithelial Neoplasias (PanINs), followed by transition to adenocarcinoma (5). More than 90% of PDAC patients have activating mutations in *KRAS* (5); however, compared with other cancer types, PDAC has a relatively low mutational load. This has impeded the discovery of therapeutic targets.

In tandem with mutational events, proteins in cancer cells are dysregulated through alterations in posttranslational modifications. Among these modifications, abnormal glycosylation has long been associated with carcinogenesis (6-8). In particular, tumor cells often display increased surface sialylation as a consequence of upregulated expression of various Golgi sialyltransferases including ST6GAL1 (9-11). ST6GAL1 is overexpressed in a wide range of malignancies (12-14), and high ST6GAL1 expression correlates with a poor prognosis in pancreatic and other cancers (15-17). ST6GAL1 adds an α2,6-linked sialic acid to select *N*-glycosylated proteins destined for the plasma membrane or secretion. Through the α2,6 sialylation of receptors such as the β1 integrin (18, 19), the Fas and TNFR1 death receptors (20, 21), and the receptor tyrosine kinases, EGFR, MET and HER2 (22-26), ST6GAL1 promotes tumor cell migration and apoptosis-resistance. ST6GAL1 also facilitates epithelial to mesenchymal transition (EMT) (23, 27) and imparts a cancer stem cell (CSC) phenotype (17, 28). The ability of ST6GAL1 to confer CSC-like characteristics may be due, in part, to its activity in upregulating the expression of stem cell-associated transcription factors such as SOX9 (17). Interestingly, as with CSCs, ST6GAL1 expression is enriched in certain nonmalignant stem cell populations and tissue niches (28, 29). Furthermore, both ST6GAL1 expression and α2,6 sialylation are increased when somatic cells are reprogrammed into induced pluripotent stem cells (iPSCs), and knockdown of ST6GAL1 hinders iPSC reprogramming (28, 30, 31). More recently, a role for ST6GAL1 in tissue regeneration has been suggested. Punch *et al*. reported that ST6GAL1 is essential for the regeneration of the gastrointestinal tract following whole-body irradiation of mice, and it was hypothesized that ST6GAL1 may function in this context to protect intestinal stem cells (32).

The de-differentiation of mature acinar cells into progenitor-like, ductal cells during the process of ADM is crucial for pancreatic regeneration following damage caused by inflammation and other injuries (2-4). Acinar cells undergoing ADM re-enter the cell cycle in order to repair the tissue. ADM-like cells typically revert to a differentiated acinar phenotype upon tissue healing (2); however, if these cells acquire oncogenic *KRAS* mutations, they undergo neoplastic transformation, leading to the generation of PanINs (2). Many studies have identified SOX9 as a critical mediator of ADM. SOX9 expression is upregulated in ADM-like cells, and genetically-engineered mouse (GEM) models have confirmed the importance of SOX9 in the formation of ADM and PanIN lesions (33, 34). In fact, Kopp *et al*. suggested that SOX9-driven ADM is a key mechanism underlying PDAC initiation (33). SOX9 expression is induced in ADM-like cells through signaling by EGFR (35). EGFR is activated within the inflammatory milieu by its ligand, TGFα, and both TGFα and EGFR are potent activators of ADM (36-39).

In view of prior evidence indicating that ST6GAL1 activates EGFR (via sialylation), stimulates SOX9 upregulation, and confers progenitor characteristics, we investigated whether ST6GAL1 facilitates the development of ADM. We first verified that ST6GAL1 is overexpressed in PDAC patient tissues, and then established a tumor-driver function for ST6GAL1 using tumor xenograft models as well as GEM models with dual expression of ST6GAL1 and oncogenic KRAS (KRAS^G12D^). Multiple *in vitro* and *in vivo* approaches were subsequently employed to show that ST6GAL1 plays a causal role in promoting ADM. Finally, we observed that acinar cells and organoid lines from mice with transgenic ST6GAL1 expression have dramatically increased activation of EGFR, highlighting a potential mechanism by which ST6GAL1-mediated receptor sialylation fuels ADM development. In the aggregate, these data suggest that ST6GAL1 may contribute to neoplasia by inducing acinar cells to adopt progenitor-like characteristics.

## Results

### ST6GAL1 is upregulated in human PDAC and promotes PDAC progression in tumor xenograft models

ST6GAL1 expression was evaluated by immunohistochemistry (IHC) in human nonmalignant pancreas and pancreatic tissues from PDAC patients using an extensively validated antibody (Fig. S1A and (14, 17, 28)). In the nonmalignant exocrine pancreas, ST6GAL1 expression was undetectable in acinar cells as well as in the great majority of ductal cells (Fig. 1A). A very small subset of ductal cells stained positively for ST6GAL1 (Fig. S1B). In contrast, strong ST6GAL1 expression was observed in the malignant cells of PDAC patients (Fig. 1A and Fig. S1B). The staining pattern was punctate and nuclear-adjacent, characteristic of Golgi localization. Golgi localization of ST6GAL1 was confirmed by co-staining for GM130 (Fig. 1B). ST6GAL1 expression was elevated in early-stage PDAC (Stage I), and levels were further increased in advanced stages (Stages III/IV) (Fig. 1C; representative IHC-stained tissues in Fig. S1C).

**Figure 1.**
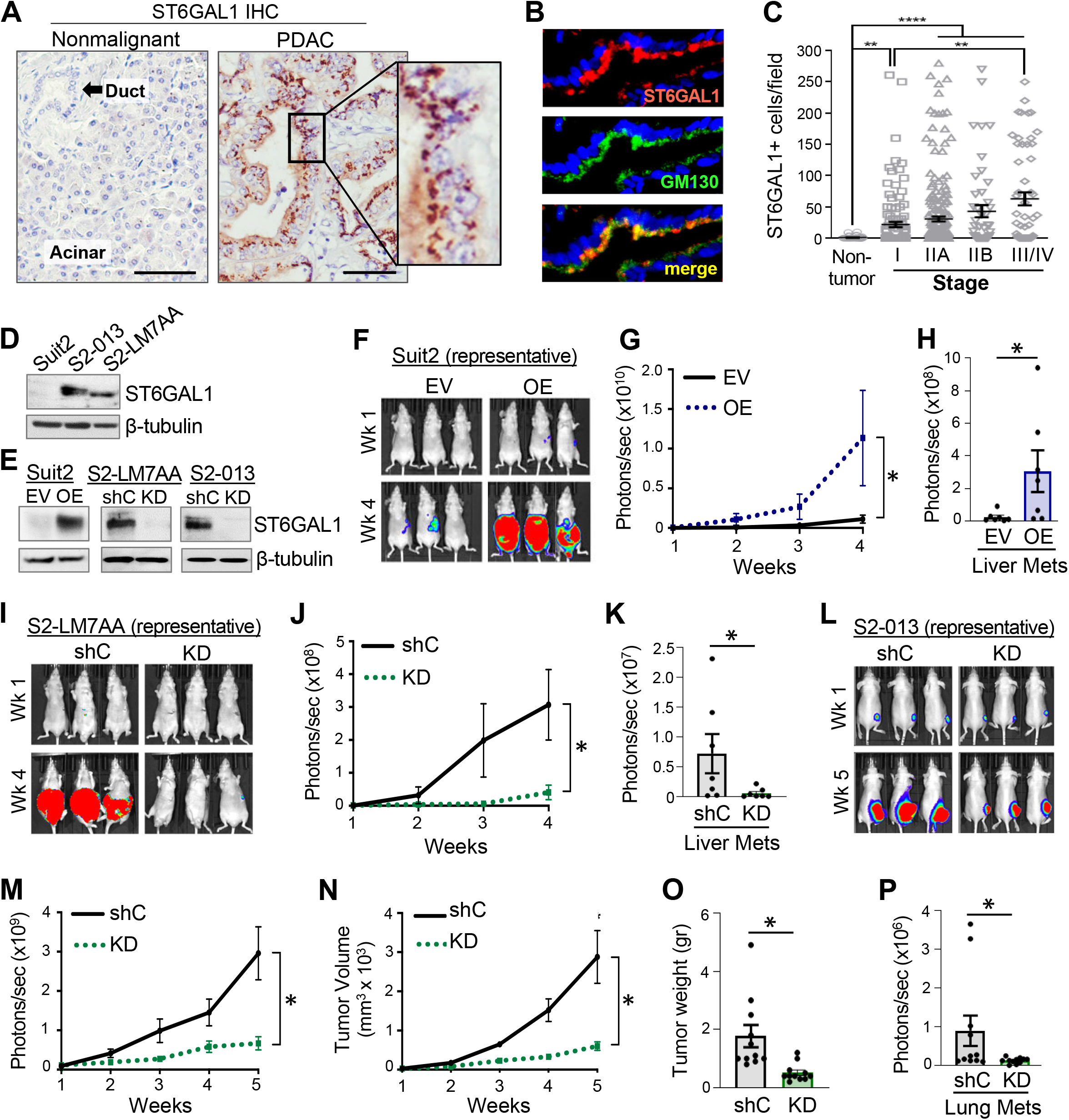
ST6GAL1 is upregulated in human PDAC and promotes tumor growth and progression in xenograft models. (A) IHC staining for ST6GAL1 in the nonmalignant human pancreas and pancreatic tissues from PDAC patients. Scale bar = 100µM. (B) PDAC specimens were co-stained for ST6GAL1 and the Golgi marker, GM130. (C) Quantification of ST6GAL1-positive cells in nonmalignant pancreata and pancreata from PDAC patients with varying stages of disease. Each symbol represents an individual patient. For: non-tumor specimens, n=48; Stage I, n=115; Stage IIA, n=216; Stage IIB, n=48; and Stage III/IV, n=55; **p<0.001; ****p<0.0001. (D) ST6GAL1 expression is enriched in the metastatic Suit2 subclones, S2-013 and S2-LM7AA, compared with the poorly-metastatic parental Suit2 line. (E) ST6GAL1 was overexpressed (OE) in Suit2 cells, with an empty vector (EV) serving as the control. ST6GAL1 was knocked-down (KD) in the metastatic S2-013 and S2-LM7AA subclones, with a non-targeting shRNA sequence serving as the control (shC). All lines represent stable, polyclonal populations. (F) Suit2 EV and OE cells were implanted into the pancreas and tumor growth monitored by BLI. Representative images are shown for mice at 1 and 4 weeks post-injection. (G) Quantification of Suit2 EV and OE tumors by BLI (n=7 mice/group). *p<0.05. (H) Livers were extracted at the endpoint from Suit2 EV and OE cohorts and imaged by BLI to detect metastases (n=7 mice/group). *p<0.05. (I) S2-LM7AA shC and KD cells were implanted into the pancreas and monitored for tumor growth by BLI. Representative images are shown for mice at 1 and 4 weeks post-injection. (J) Quantification of tumor growth in mice implanted with S2-LM7AA shC and KD cells (n=7 mice/group) *p<0.05. (K) Quantification of metastatic tumors by BLI of livers extracted at the endpoint from S2-LM7AA shC and KD cohorts (n=7 mice/group). *p<0.05. (L) Representative images of mice at 1 and 5 weeks following injection of S2-013 shC and KD cells into the flank. (M) Quantification by BLI of flank tumors formed from S2-013 shC and KD cells (n=11 mice/group). *p<0.05. (N) Tumor volume was calculated by caliper measurements (n=11 mice/group). *p<0.05. (O) Weights of tumors extracted at the endpoint (n=11 mice/group).*p<0.05. (P) Lungs extracted at the endpoint were evaluated by BLI to quantify metastatic burden (n=11 mice/group). *p<0.05

To interrogate a putative pro-tumorigenic function for ST6GAL1, tumor xenograft experiments were conducted using the human Suit2 PDAC cell line, and two metastatic Suit2 subclones. Although ST6GAL1 is robustly expressed in the majority of human PDAC cell lines, Suit2 cells are unusual in that they have low levels of ST6GAL1 and limited metastatic potential (23). Other groups have used iterative *in vivo* selection methods to isolate Suit2 subclones with enhanced metastatic capability (40, 41). Two such subclones, S2-LM7AA and S2-013, have greatly upregulated ST6GAL1 relative to parental Suit2 cells (Fig. 1D), suggesting that ST6GAL1 expression is selected for during metastatic progression. ST6GAL1 was overexpressed (OE) in Suit2 parental cells, or knocked-down (KD) in S2-LM7AA and S2-013 cells (Fig. 1E), and the expected changes in α2,6 surface sialylation were confirmed by staining cells with the SNA lectin (Fig. S2A). Suit2 OE cells or empty vector (EV) controls were implanted into the pancreas, and tumor growth monitored by bioluminescence imaging (BLI). As shown in Fig. 1F-G, significantly increased tumor growth was observed for OE cells relative to EV cells, and imaging of livers extracted at the endpoint revealed a greater metastatic burden for the OE cohort (Fig. 1H, representative images in Fig. S2B). ST6GAL1 activity similarly promoted tumor growth in the S2-LM7AA cell model, evidenced by the reduced growth of pancreatic tumors (Fig. 1I-J) and decreased metastasis (Fig. 1K) in the KD cohort compared with shC controls. ST6GAL1 KD in the S2-013 cell model also impaired tumor growth; however, for this line, cells were injected into the flank and monitored for metastasis to the lungs, as in prior studies (41). Mice injected with S2-013 KD cells had reduced growth of flank tumors as indicated by BLI (Fig. 1L-M), tumor volume (Fig. 1N), and tumor weight (Fig. 1O). Reduced metastasis was also observed in the S2-013 KD cohort relative to shC controls (Fig. 1P). The growth of primary and metastatic tumors for the three lines was confirmed by histology (Fig. S2C-D). Together, xenograft experiments using the isogenic Suit2 cell series provide strong evidence that ST6GAL1 has pro-tumorigenic activity.

### ST6GAL1 overexpression in mice expressing KRAS^G12D^ promotes accelerated PDAC progression and mortality

To model the upregulation of ST6GAL1 observed in human PDAC, we generated a transgenic mouse line with conditional expression of human ST6GAL1 in the pancreas. C57BL/6 mice were engineered with an *LSL-ST6GAL1* transgene inserted into the Rosa26 locus. These mice were crossed to the *Pdx1-Cre* line to direct pancreatic expression of ST6GAL1 (*Pdx1-Cre; LSL-ST6GAL1*, abbreviated as “SC” mice). IHC confirmed strong ST6GAL1 expression in the pancreatic acinar cells of SC mice (Fig. 2A), whereas no detectable ST6GAL1 was observed in the acinar cells of littermate controls harboring the *ST6GAL1* transgene, but lacking Cre (*LSL-ST6GAL1*, hereafter referred to as wild-type, “WT”, for brevity). SNA staining of cells dissociated from WT and SC pancreata revealed increased surface α2,6 sialylation of SC cells (Fig. 2B). Necropsy performed on neonatal SC mice indicated that there were no developmental abnormalities in the pancreas or other organs (Fig. S3A-B). To study PDAC development, we used the “KC” model (42), in which *Pdx1-Cre* drives KRAS^G12D^ expression in the pancreas (*Pdx1-Cre; LSL-Kras*^*G12D*^). The various lines were crossed to generate “KSC” mice, which have expression of both KRAS^G12D^ and ST6GAL1 (*Pdx1-Cre; LSL-ST6GAL1; LSL-Kras*^*G12D*^). No pancreatic abnormalities were observed in neonatal KC or KSC mice, although PanINs were detected at 8 weeks in both models (Fig. S3C-D). Notably, endogenous ST6GAL1 was strongly expressed in the PanIN lesions of KC mice (Fig. 2C, arrow), illustrating an upregulation of ST6GAL1 in early stages of neoplasia. In KSC mice, ST6GAL1 was also expressed in the PanINs, however, unlike KC mice, the adjacent, normal acinar cells displayed strong ST6GAL1 staining (Fig. 2C, arrowhead), reflecting expression of the *ST6GAL1* transgene.

**Figure 2.**
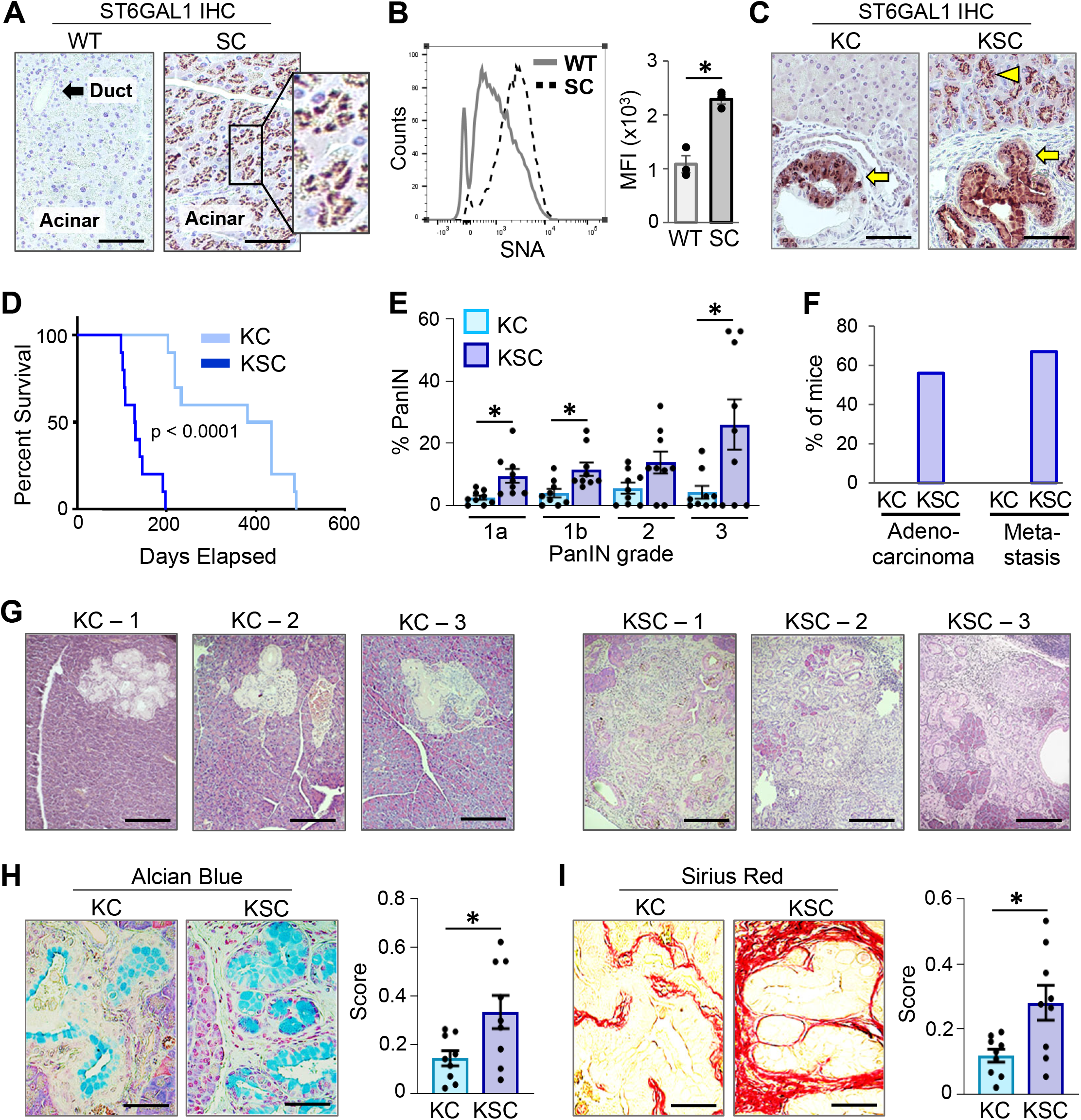
KSC mice exhibit accelerated PDAC progression and mortality compared to KC mice. (A) ST6GAL1 expression in the pancreas of SC mice (with *Pdx1-Cre*-driven expression of *ST6GAL1*), and “WT” mice (littermate controls expressing the *LSL-ST6GAL1* transgene, but not *Pdx1-Cre*). In SC mice, ST6GAL1 staining in acinar cells appears in a punctate pattern characteristic of Golgi localization. Scale bar = 100µM. (B) Cells dissociated from WT and SC pancreata were stained with the SNA lectin to detect surface α2,6 sialylation. Left panel shows a representative flow cytometry experiment; right panel depicts Mean Fluorescent Intensity (MFI) from 3 mice per genotype. *p<0.05. (C) IHC staining of pancreata from KC and KSC mice reveals strong ST6GAL1 expression in the neoplastic PanINs (arrows). Conversely, staining in the adjacent, normal-appearing acinar cells of KC vs. KSC mice is starkly different. No detectable ST6GAL1 is noted in normal KC acinar cells, whereas strong staining is observed in normal KSC acinar cells (arrowhead), reflecting expression of the *ST6GAL1* transgene. Scale bar = 100µM. (D) Kaplan-Meier survival analysis indicates a median survival of 4.3 months for KSC mice and 13.6 months for KC mice. (n=10 mice/group) *p<0.0001. (E) H&E-stained pancreata from 20 week-old KC and KSC mice were examined for percentage of overall tissue area represented by PanINs of varying grades (n=9 mice/group). Tissues were evaluated under blinded conditions by a board-certified pathologist, Dr. Isam Eltoum. *p<0.05. (F) Tissues from the 20 week-old cohorts were evaluated for the presence of adenocarcinoma in the pancreas, or metastatic tumors in the liver, lungs, intestines, or peritoneum (n=9 mice/group). (G) Representative pancreatic tissues showing more advanced disease in KSC mice. (KC 1-3 represent 3 individual KC mice; KSC 1-3 represent 3 individual KSC mice). Scale bar = 200µM. H) Alcian Blue staining for mucinous tumor cells in the 20 week-old KC and KSC cohorts (n=9 mice/group). Staining was quantified by stereological analysis. *p<0.05. Scale bar = 100µM. (I) Sirius Red staining for collagen deposition in the pancreata of 20 week-old KC and KSC mice (n=9 mice/group). Staining was quantified by stereological analysis. *p<0.05. Scale bar = 100µM.

A Kaplan-Meier survival analysis revealed dramatically accelerated mortality for KSC mice (Fig. 2D), evidenced by a median survival of 4.3 months, as compared with 13.6 months for KC mice. The 13.6 month survival time for KC mice is comparable to that reported by others (43). Analyses of tissues harvested from the survival cohort confirmed pancreatic malignancy as the cause of mortality for KC and KSC mice (Fig. S3E).

### ST6GAL1 promotes PanIN formation, adenocarcinoma and metastasis

To compare pathogenesis in age-matched KC and KSC mice, we generated 20 week-old cohorts and analyzed tissues for the presence of PanINs, adenocarcinoma, and distal metastases. In the KC cohort, 2/9 mice had no apparent pancreatic abnormalities, whereas all 9 KSC mice displayed PanINs. KSC mice had more extensive overall PanIN development than KC mice, and a particularly prominent increase in advanced grade PanIN3 (carcinoma in situ) was noted (Fig. 2E). Additionally, 5/9 (56%) of the KSC cohort presented with adenocarcinoma, and 6/9 KSC mice (67%) had metastases to the liver, lungs and/or other sites (Fig. 2F, representative images in Fig. S3F). None of the KC mice developed adenocarcinoma or metastatic disease at the 20 week time point. Images of H&E-stained pancreata revealed more advanced disease and fibrosis in 20 week-old KSC mice (Fig. 2G). Staining of tissues with Alcian Blue (Fig. 2H), and Sirius Red (Fig. 2I) confirmed that KSC mice had a greater abundance of mucinous tumor cells and more extensive desmoplasia than KC mice.

### RNA-Seq analyses of GEM pancreata show that ST6GAL1 activity upregulates stem and cancer-associated gene networks, promotes a pancreatic ductal cell program, and induces activation of EGFR

Given ST6GAL1’s known role in promoting progenitor-like characteristics, we hypothesized that ectopic expression of ST6GAL1 in acinar cells may contribute to tumorigenesis by upregulating stem and ductal gene networks that facilitate both ADM and neoplastic transformation. To obtain an unbiased view of the effects of ST6GAL1 on pancreatic cell phenotype, RNA-Seq was conducted on pancreata from 20 week-old WT and SC mice. At 20 weeks of age, there is no evidence of malignancy or other abnormalities in SC mice. In the normal pancreas, acinar cells comprise 80-90% of the total mass, therefore acinar transcripts dominate the total mRNA pool (44). We used Ingenuity Pathway Analysis (IPA) to identify the top 5 Associated Network Functions that differed between SC and WT mice (Fig. 3A). Developmental processes were heavily represented, exemplified by embryonic development, organismal development, and organ development. Gene Set Enrichment Analyses (GSEA) revealed an enrichment in developmental and stemness pathways in SC cells including ESC pluripotency, Wnt, Notch, and Hedgehog, and their associated downstream signaling nodes including β-catenin (Wnt pathway) and Hes-Hey (Notch pathway) (Fig. 3B and Table S1). The Wnt, Notch, and Hedgehog pathways play pivotal roles in pancreatic exocrine development, and PDAC is marked by re-activation of these pathways (45). Importantly, SC cells displayed an enrichment in a pancreatic ductal cell program (Fig. 3B), and a heatmap of this dataset (Fig. 3C) indicated increased expression of canonical ductal markers in SC cells including Sox9, Onecut2, Muc1, Hnf1b, Litaf, and Slc4a4. Additionally, TGFα and EGFR levels were increased, which is noteworthy given the prominent role of TGFα/EGFR signaling in driving ADM (38, 46, 47). Nectin4 expression was also elevated in SC mice. EGFR activation induces the de-differentiation of acinar cells into Nectin-positive progenitors that serve as a transitional population between differentiated acinar cells and fully transdifferentiated ADM-like cells (47). GSEA confirmed that EGFR and other ERBB signaling networks were activated in SC mice (Fig. 3D). Several other receptor tyrosine kinases were also activated in SC cells such as MET (Fig. 3D and Table S1). Both MET and EGFR are upregulated in PanINs and PDAC (5). Of particular interest, numerous cancer networks were elevated in SC mice including pancreatic (Fig. 3B), breast, renal, lung, prostate and other cancers (Table S1). The activation of cancer-associated networks in SC cells aligns with the concept that ST6GAL1 upregulation induces molecular signaling events and transcriptomic changes that may predispose cells to neoplastic transformation.

**Figure 3.**
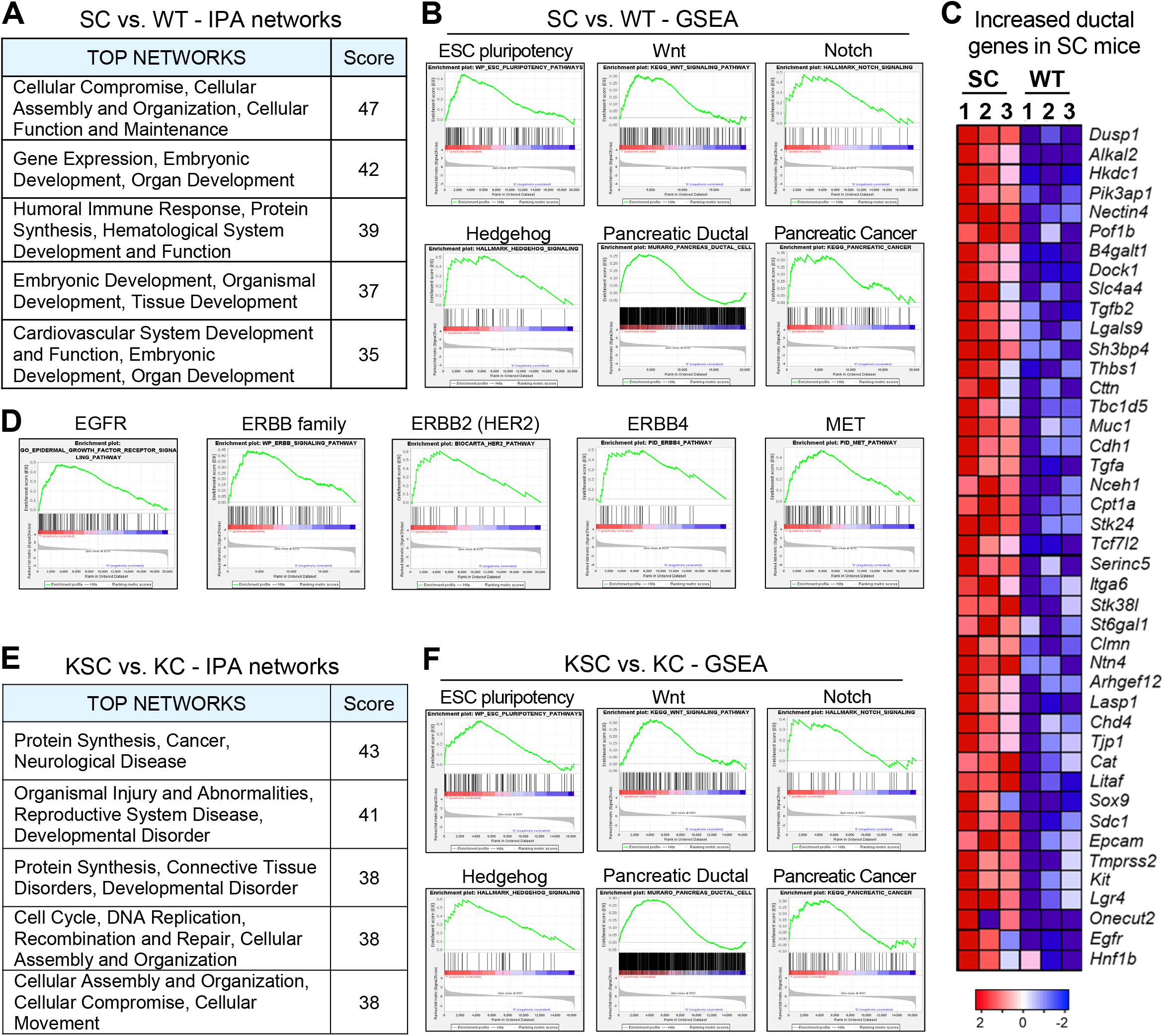
RNA-Seq experiments reveal that ST6GAL1 activity promotes a stem and ductal phenotype, and enhances activation of EGFR and other ERBB family members. (A) Ingenuity Pathway Analysis (IPA) of RNA-Seq data from 20-week old SC and WT mice. The 5 top-scoring networks altered in SC versus WT mice are shown (n=3 mice/group). (B) GSEA comparing 20-week old SC and WT mice indicates that SC mice have an upregulation in stemness networks (ESC pluripotency, Notch, Wnt, Hedgehog) as well as networks associated with a pancreatic ductal cell program and pancreatic cancer. NES and FDR values are as follows: <u>ESC pluripotency</u>: NES=2.04, FDR=0.011; <u>Wnt:</u> NES=1.48, FDR=0.103; <u>Notch:</u> NES=1.70, FDR=0.008; <u>Hedgehog:</u> NES=1.89, FDR=0.003; <u>Ductal:</u> NES=1.59, FDR=0.018; <u>Pancreatic cancer</u>: NES=1.44, FDR=0.135 (C) Heatmap of select genes from the pancreatic ductal network upregulated in SC mice. SC1-3 represent 3 individual SC mice; WT1-3 represent 3 individual WT mice. (D) GSEA reveals activation of EGFR and other receptor tyrosine kinases in SC mice. <u>EGFR:</u> NES=2.22, FDR=0.003; <u>ERBB family:</u> NES=1.96, FDR=0.015; <u>ERBB2:</u> NES=1.96, FDR=0.049; <u>ERBB4:</u> NES=1.78, FDR=0.023; <u>MET:</u> NES=2.04, FDR=0.010. (E) IPA of RNA-Seq data from 20-week old KSC and KC mice depicting the 5 top-scoring networks (n=3 mice/genotype). (F) GSEA comparing 20 week-old KC and KSC mice indicates that KSC mice have an upregulation in stemness-associated networks (ESC pluripotency, Notch, Wnt and Hedgehog) and networks associated with a ductal phenotype and pancreatic cancer. NES and FDR values are as follows: <u>ESC pluripotency:</u> NES=1.84, FDR=0.033; <u>Wnt:</u> NES=1.49, FDR=0.123; <u>Notch:</u> NES=1.38, FDR=0.077; <u>Hedgehog:</u> NES=2.09, FDR<0.0005; <u>Ductal:</u> NES=1.65, FDR=0.019; <u>Pancreatic cancer:</u> NES=1.26, FDR=0.280.

We next compared 20 week-old KC and KSC mice. At this age, most KSC mice have advanced malignancy with extensive desmoplasia, whereas KC mice harbor mostly early-stage PanINs. Thus, many, if not most, of the observed changes in gene expression are likely to be secondary to these divergent stages in disease progression. IPA analysis of KSC versus KC pancreata indicated that the top 5 Associated Network Functions related to developmental disorders, organismal injury and abnormalities, cancer, protein synthesis, and cellular compromise, assembly and movement (Fig. 3E). GSEA revealed that, compared to KC mice, KSC mice had an enrichment in: (i) stem cell networks (ESC pluripotency, Wnt, Notch, and Hedgehog); (ii) a pancreatic ductal cell program; (iii) cancer-associated networks; and (iv) EGFR, ERBB2/HER2, and MET signaling pathways (Fig. 3F and Table S2).

### Nonmalignant pancreatic acinar cells of mice with ectopic ST6GAL1 expression exhibit an upregulation in SOX9 and other ductal markers

To confirm activation of a ductal cell program in SC acinar cells, pancreata from WT and SC mice were IHC stained for SOX9 (a nuclear-localized transcription factor). SOX9 was not detected in the acinar cells of WT mice, in stark contrast with the large number of SOX9-positive acinar cells in SC pancreata (Fig. 4A-B). In KC and KSC mice, SOX9 was highly expressed in the PanIN lesions (“P”) (Fig. 4C), consistent with the well-known upregulation of SOX9 during malignant transformation (33). However, similar to SC mice, KSC mice displayed extensive SOX9 expression in adjacent, morphologically-normal acinar cells, whereas few of the normal-appearing acinar cells of KC mice had SOX9 expression (Fig. 4D). We also evaluated other classic ductal markers upregulated during both ADM and pancreatic neoplasia, specifically, cytokeratins 8 and 19 (KRT 8/19). Both of these cytokeratins were pervasively expressed by the acinar cells of SC mice, but not WT acinar cells (Fig. 4E-F). These data indicate that ectopic expression of ST6GAL1 in acinar cells is sufficient to induce the upregulation of ductal genes. Of note, the acinar cells of SC mice did not display any obvious morphological differences from WT acinar cells. Prior studies have suggested that mature acinar cells de-differentiate through a series of stages including an initial transition into an “acinar-like” cell with a normal morphology, but increased expression of progenitor genes and decreased levels of some zymogens (4). This is followed by transition into a progenitor-like cell, and then finally, transdifferentiation into ductal-like cells with sharply reduced acinar markers and a tubular morphology characteristic of ADM (4).

**Figure 4.**
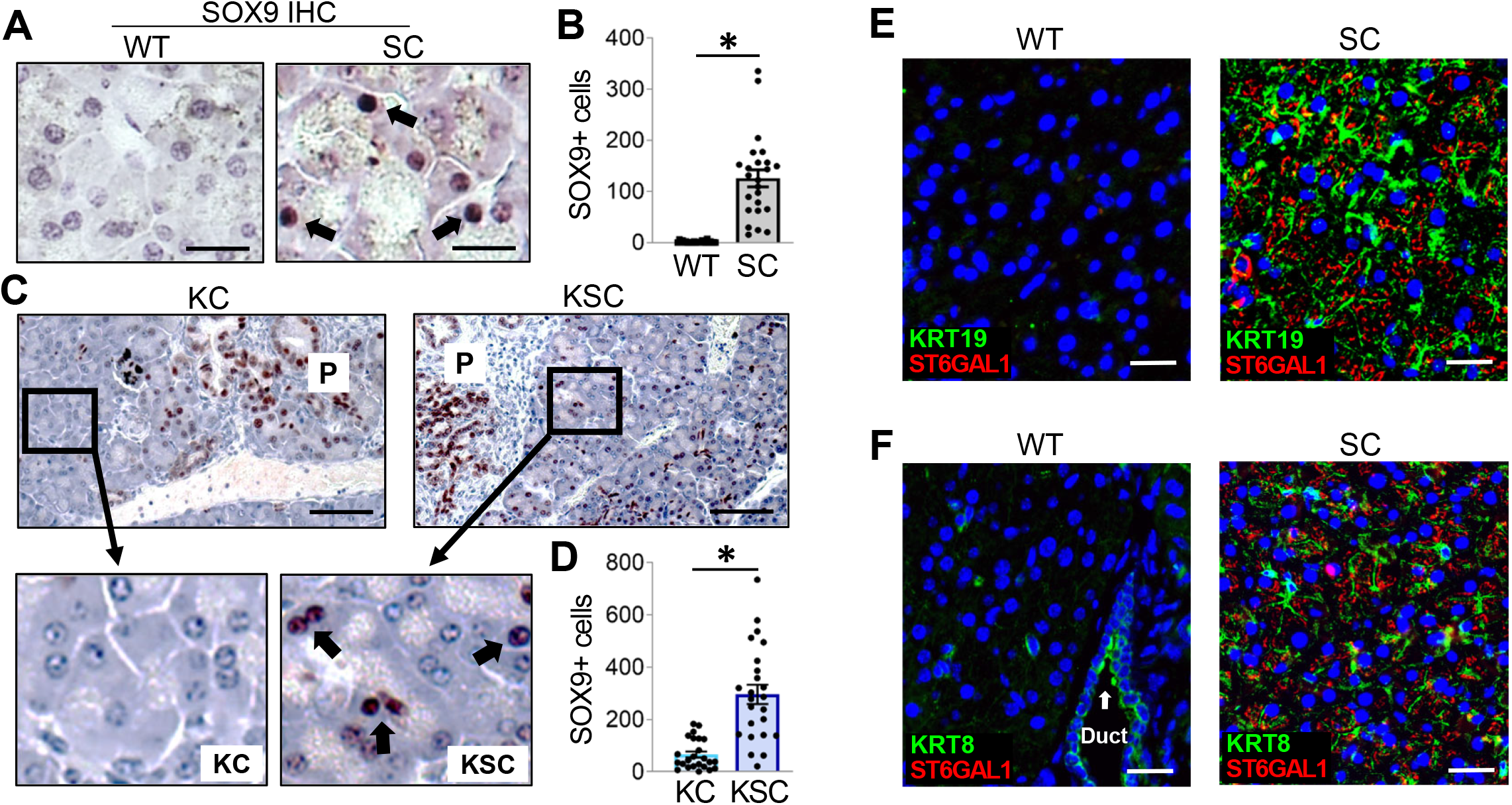
Acinar cells from mice with ectopic expression of ST6GAL1 have upregulated expression of SOX9 and other ductal markers. (A) IHC staining of pancreata from 20-week old mice reveals SOX9 expression (arrows) in the acinar cells of SC, but not WT, mice. Scale bar = 20µM. (B) SOX9-expressing acinar cells were quantified from IHC-stained pancreata from 20 week-old SC and WT mice (n=8 mice/group, with 3 tissue sections evaluated per mouse). *p<0.05. (C) IHC staining for SOX9 in pancreata from 20 week-old KC and KSC mice. “P” denotes PanINs, which are positive for SOX9. Scale bar = 100µM. Insets show upregulation of SOX9 (arrows) in the nonmalignant, normal-appearing acinar cells of KSC, but not KC, mice. (D) Quantification of SOX9-positive cells in the normal-appearing acinar cells of 20 week-old KC and KSC mice (n=8 mice/group, with 3 tissue sections evaluated per mouse). *p<0.05. (E) WT and SC pancreata were stained for ST6GAL1 (red) and cytokeratin 19 (KRT19, green). Nuclei were counterstained with Hoechst (blue). Scale bar = 25µM. (F) WT and SC pancreata were stained for ST6GAL1 (red) and cytokeratin 8 (KRT8, green), and counterstained with Hoechst (blue). Note that ductal cells (but not acinar cells) in WT mice have high expression of KRT8. Scale bar = 25µM.

### Pancreatic organoids from mice with ectopic ST6GAL1 expression have enhanced growth and increased expression of stem and ductal genes

Organoid lines were generated from the GEM models. Organoids initiate from progenitor-like cells, and maintain progenitor properties during propagation in media containing stem cell factors (48). However, placing dissociated organoid cells into 2D confluent, monolayer culture in media with reduced stem cell factors causes cells to undergo differentiation (29, 49). Upon placement of cells from WT organoids into monolayer culture (ML), endogenous *St6gal1* was downregulated, similar to the hallmark stemness genes, *Lgr5* and *Axin2* (Fig. 5A).

**Figure 5.**
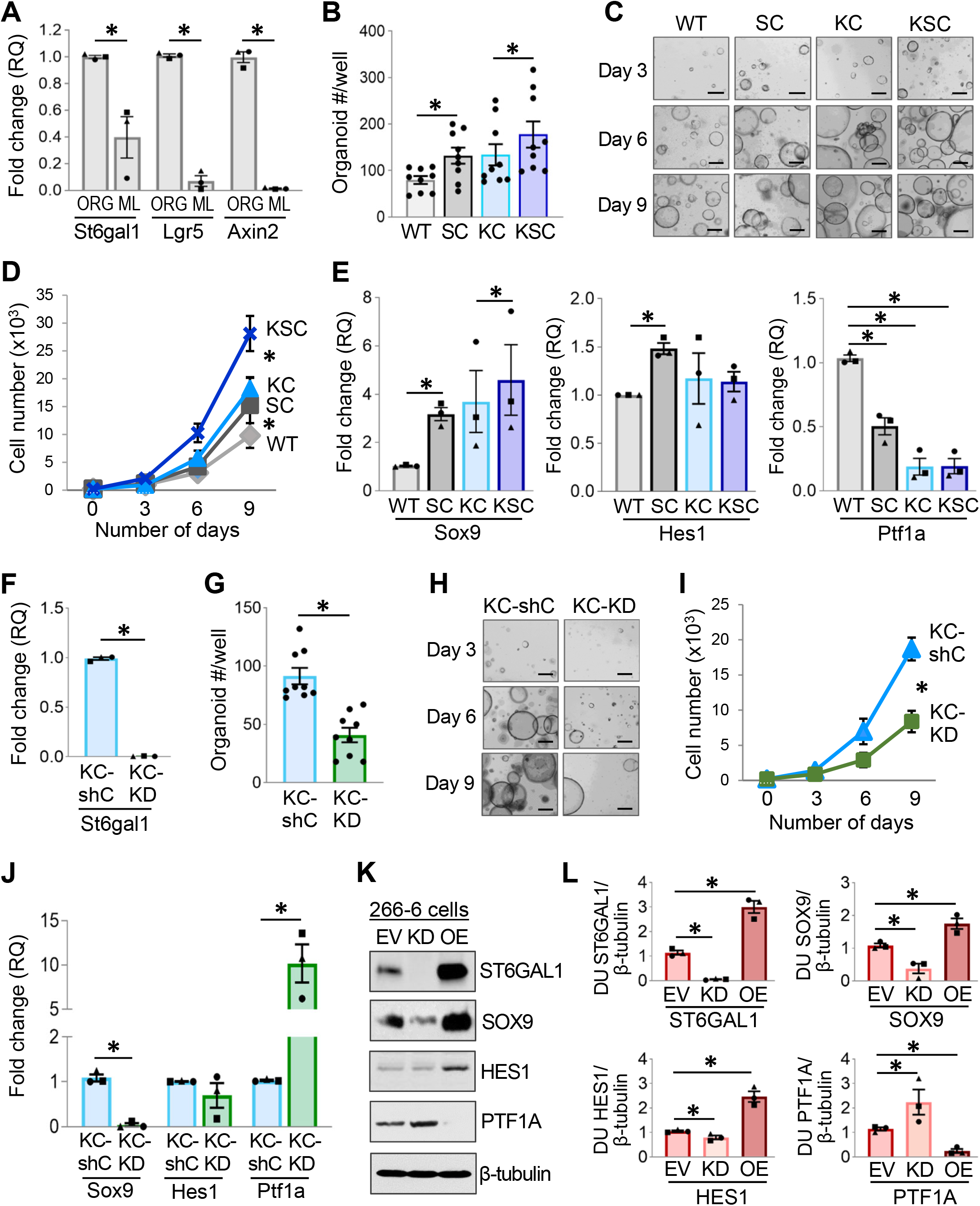
ST6GAL1 activity enhances the growth of GEM-derived organoids and promotes a stem/ductal gene program in both organoids and the 266-6 ADM cell model. (A) Cells dissociated from WT organoids were seeded into either organoid (ORG) culture, which maintains stemness properties, or monolayer (ML) culture in media with reduced stem cell factors. qRT-PCR analyses revealed that, similar to the stemness genes *Axin2* and *Lgr5*, endogenous *St6gal1* mRNA expression was downregulated in ML cultures. Three independent experiments were conducted, with each experiment performed in triplicate. *p<0.05. (B) Cells were dissociated from organoid lines, and 2,000 cells seeded into fresh organoid culture. The number of organoids formed at Day 3 was enumerated. Three independent experiments were performed, with 3 wells per genotype counted in each experiment. *p<0.05. (C) Cells were dissociated from the organoid lines, seeded into fresh organoid culture, and organoid growth was monitored over time. Representative images of organoid cultures are shown. Scale bar = 100µM. (D) At Days 3, 6, and 9 following the seeding of 2,000 organoid-derived cells into fresh organoid culture, organoids were dissociated, and the total number of cells was quantified. Three independent experiments were conducted. *p<0.05. (E) mRNA levels of *Sox9, Hes1* and *Ptf1a* were quantified in the organoids by qRT-PCR. Three independent experiments were conducted, with each experiment performed in triplicate. *p<0.05. (F) KC organoids were transduced with lentivirus encoding *St6gal1* shRNA (KC-KD) or a control shRNA vector (KC-shC). Knockdown of *St6gal1* was verified by qRT-PCR. Three independent experiments were conducted, with each experiment performed in triplicate. *p<0.05. (G) Cells were dissociated from the KC-shC and KC-KD organoids, and 2,000 cells seeded into fresh organoid culture. The number of organoids formed at Day 3 was enumerated. Three independent experiments were conducted, with 3 wells counted per genotype in each experiment. *p<0.05. (H) Representative images of the KC-shC and KC-KD cultures are shown. Scale bar = 100µM. (I) At Days 3, 6, and 9 after seeding 2,000 organoid-derived cells into fresh organoid culture, organoids were dissociated, and the total number of cells was quantified. Three independent experiments were conducted. *p<0.05. (J) mRNA levels of *Sox9, Hes1* and *Ptf1a* were quantified by qRT-PCR in KC-shC and KC-KD organoids. Three independent experiments were conducted, with each performed in triplicate. *p<0.05. (K) ST6GAL1 was overexpressed (OE) or knocked-down (KD) in the 266-6 pancreatic cancer cell line, a common ADM model. An empty-vector (EV) construct served as the control. The expression of SOX9, HES1 and PTF1A was measured by immunoblotting. (L) Densitometry was conducted on immunoblots from 3 independently-generated 266-6 cell lysates. Densitometric Unit (DU) measurements for target proteins were normalized to DUs for β-tubulin. *p<0.05.

We next compared the 4 GEM models for organoid-forming capability. The ability of dissociated, single cells to form organoids is an indicator of self-renewal potential. At 3 days following cell seeding into organoid culture, a greater number of organoids developed from SC compared with WT cells, and KSC compared with KC cells (Fig. 5B). Additionally, organoid growth over a 9-day interval was enhanced in the SC and KSC organoids relative to the WT and KC organoids, respectively (Fig. 5C-D). We then assessed the expression of the ADM-associated stem/ductal genes, *Sox9* and *Hes1*, as well as the differentiated acinar marker, *Ptf1a*. Compared with WT organoids, SC organoids had significantly increased expression of *Sox9* and *Hes1*, but reduced expression of *Ptf1a* (Fig. 5E). Interestingly, lineage tracing experiments have shown that only the SOX9-expressing ductal-like cells are capable of long-term clonal expansion from single cells, whereas PTF1A-positive cells fail to expand (50). Comparing KC and KSC organoids, *Sox9* was increased in the KSC organoids, but there were no differences in *Hes1* or *Ptf1a* expression (Fig. 5E).

### Knockdown of ST6GAL1 in KC organoids suppresses organoid growth and ductal gene expression, while increasing acinar gene expression

To determine whether ST6GAL1 was important for maintaining progenitor characteristics in cells expressing KRAS^G12D^, ST6GAL1 expression was knocked-down in KC organoids using shRNA (KC-KD). Downregulation of *St6gal1* was confirmed by qRT-PCR (Fig. 5F). Suppression of ST6GAL1 expression in KC organoids inhibited initial organoid formation (Fig. 5G) and organoid growth over time (Fig. 5H-I). Moreover, cells with ST6GAL1 KD had decreased expression of *Sox9*, and increased levels of *Ptf1a* (Fig. 5J), suggesting that ST6GAL1 KD reverted KRAS^G12D^-expressing cells to a more differentiated, acinar phenotype. Thus, for both nonmalignant and neoplastic cells, ST6GAL1 activity was important for the initiation and growth of organoids, and also contributed to the maintenance of a ductal gene program.

### ST6GAL1 promotes the expression of ductal markers in the 266-6 ADM cell model

The acinar-like 266-6 pancreatic cancer cell line is a well-established cell model for studying ADM (51). Accordingly, 266-6 cells with ST6GAL1 overexpression (OE) or knockdown (KD) were evaluated for SOX9, HES1 and PTF1A expression by immunoblotting. Relative to EV controls, cells with ST6GAL1 KD had markedly reduced SOX9, and modestly increased PTF1A, whereas cells with ST6GAL1 OE had increased SOX9 and HES1 expression, but sharply reduced PTF1A (Fig. 5K-L).

### ST6GAL1 activity enhances inflammation-induced ADM

Pancreatitis is a potent inducer of ADM; we therefore examined ST6GAL1 activity in the cerulein-stimulated pancreatitis model (51). Upon injection of WT mice with cerulein, acinar cells adopted a tubular-like morphology characteristic of inflammatory damage and ADM (Fig. 6A). Additionally, acinar cells (identified by Amylase expression) from the cerulein-treated cohort co-expressed SOX9, a defining feature of ADM (Fig. 6B, note that the intensity of Amylase staining is reduced in ADM-like cells, consistent with the known downregulation of amylase genes in acinar cells undergoing ADM). We then probed for expression of endogenous ST6GAL1 in WT mice treated with cerulein. As shown in Fig. 6C, ST6GAL1 was not detected in the acinar cells of saline-treated controls, whereas pronounced ST6GAL1 expression was observed in the ADM-like cells (co-expressing Amylase and SOX9) of cerulein-treated mice. These data show that the upregulation of endogenous ST6GAL1 in acinar cells is a novel marker for cells undergoing inflammation-induced ADM.

**Figure 6.**
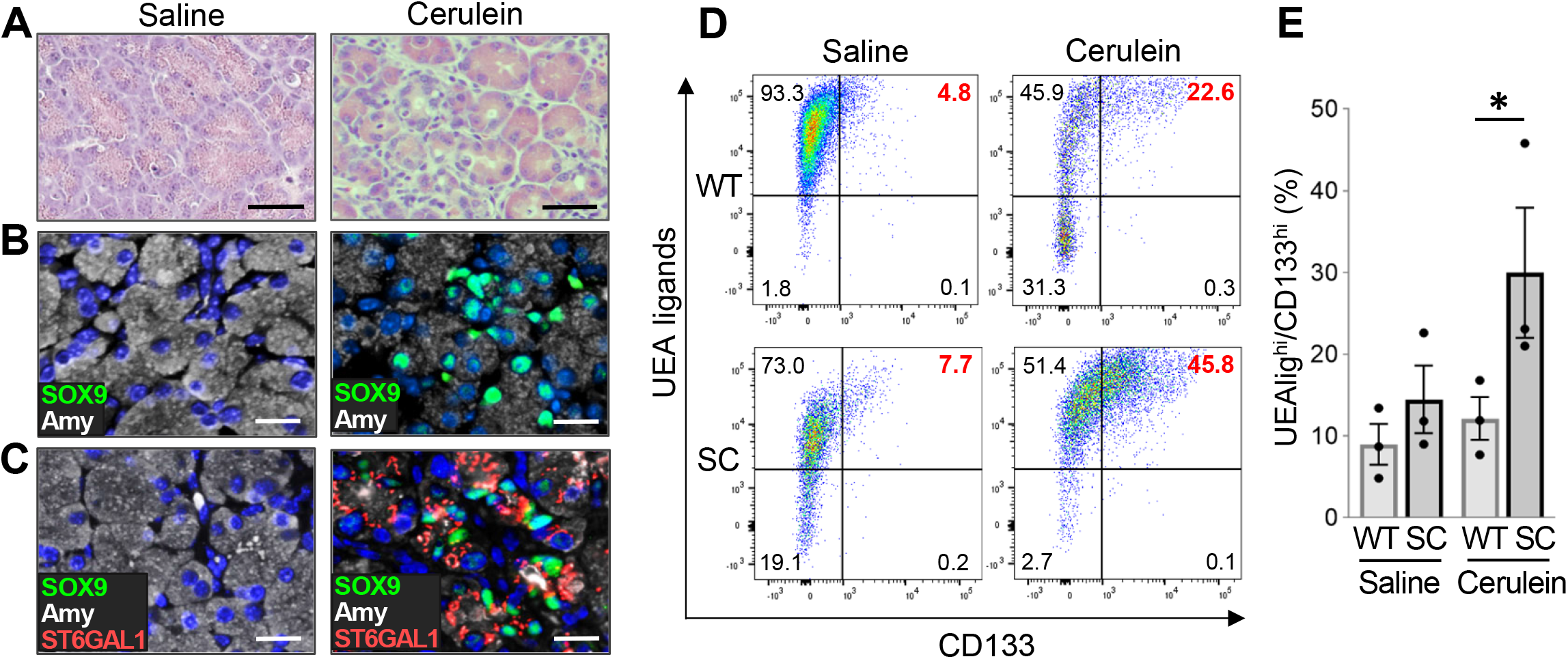
ST6GAL1 activity promotes ADM in the cerulein-induced pancreatitis model. (A) WT mice were injected i.p. with cerulein to induce pancreatitis, or with a saline control. Cerulein-induced damage to the pancreas was confirmed by H&E. Scale bar = 50µM. (B) Pancreata from WT mice injected with saline or cerulein were evaluated for ADM by co-staining for SOX9 (green) and Amylase (Amy, white). Tissues were counterstained with Hoechst (blue). Amylase and SOX9 co-expression was noted in acinar cells from cerulein-treated, but not saline-treated, mice. Scale bar = 25µM. (C) Pancreata from WT mice injected with saline or cerulein were evaluated for co-expression of SOX9 (green), Amylase (white) and ST6GAL1 (red) in ADM-like cells. Nuclei were stained with Hoechst (blue). Cells co-expressing ST6GAL1, SOX9 and Amylase were identified in cerulein-treated, but not saline-treated, mice. Scale bar = 25µM. (D) Cells were dissociated from the pancreata of WT and SC mice treated with saline or cerulein. Flow cytometry was conducted on cells stained with UEA lectin (acinar marker), anti-CD133 (ductal marker), anti-EpCAM (epithelial marker), anti-CD45 (immune marker) and Aqua live/dead stain (marker for non-viable cells). UEA and anti-CD133 staining was quantified in viable, singlet epithelial cells (EpCAM positive, CD45 negative, Aqua dye negative). Cells undergoing ADM are positive for both UEA and anti-CD133 staining. A representative experiment is shown. (E) Quantification of cells undergoing ADM (UEAlig^hi^/CD133^hi^) in cerulein-treated mice or saline-treated controls (3 mice/group). *p<0.05.

We then quantified the number of ADM-like cells in cerulein-treated WT and SC mice by staining dissociated pancreatic cells for surface markers of ADM. Differentiated acinar cells express glycan ligands recognized by the UEA lectin, however, they lack expression of CD133. Conversely, ductal cells express CD133, but not UEA ligands. In pancreatitis tissues, the co-expression of UEA ligands with high surface CD133 (UEA-ligand^hi^/CD133^hi^) is an indicator of ADM (52). As shown in Fig. 6D-E, cerulein-treated SC mice had significantly more UEA-ligand^hi^/CD133^hi^ cells than WT mice, demonstrating that ectopic expression of ST6GAL1 in acinar cells actively promotes ADM within an inflammatory milieu.

### EGFR is activated in acinar cells with ectopic ST6GAL1 expression

To investigate the role of receptor sialylation in impelling acinar cells to adopt ductal-like characteristics, we focused on the activation state of EGFR. ST6GAL1-mediated sialylation of EGFR is a strong activator of this receptor (22-24), and EGFR is one of the most potent drivers of ADM. Phosphorylated (activated) EGFR was not detected in the acinar cells of WT mice, although a low level of staining was apparent in some of the ducts (Fig. 7A, arrow). Strikingly, most of the acinar cells of SC mice displayed robust activation of EGFR. Similar differences were noted when comparing KC and KSC mice. The morphologically normal acinar cells of KC mice lacked p-EGFR expression, whereas the normal acinar cells of KSC mice had high levels of p-EGFR. On the other hand, EGFR was strongly activated in the PanIN lesions of both KC and KSC mice, consistent with an extensive literature (Fig. 7B). Finally, we examined EGFR activation in organoids by immunoblotting. As shown in Fig. 7C-E, pronounced activation of EGFR was detected in SC and KSC organoids, but not in WT and KC organoids. Collectively, results in Fig. 7 suggest that the sialylation-dependent activation of EGFR may constitute an important mechanism by which ST6GAL1 activity reprograms acinar cells into a more ductal, progenitor-like state.

**Figure 7:**
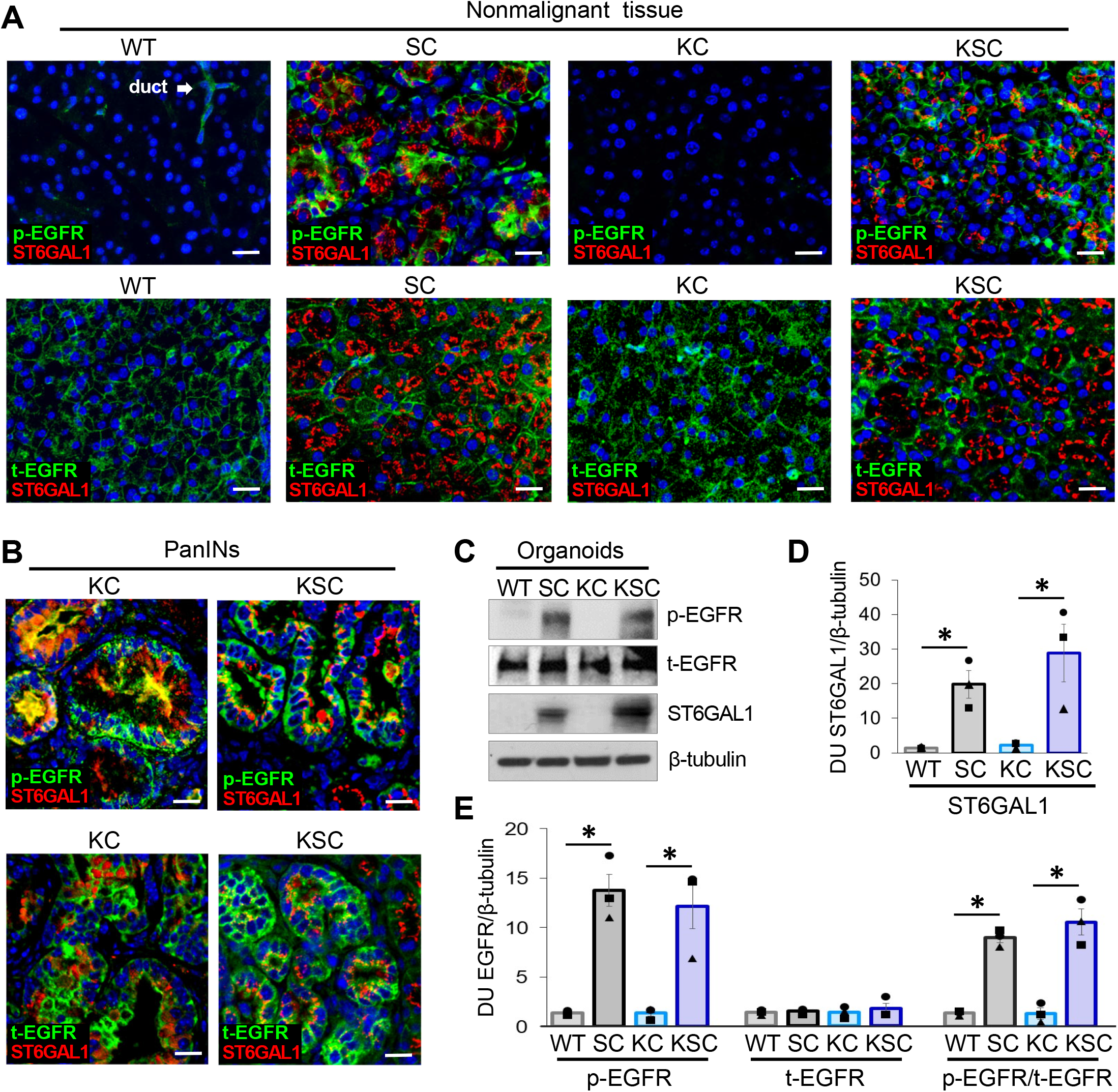
EGFR is activated in pancreatic tissues and organoids from mice with ectopic expression of ST6GAL1. (A) Upper panels: pancreata from WT and SC mice, as well as adjacent, nonmalignant tissues from KC and KSC mice, were stained for ST6GAL1 (red) and phosphorylated EGFR (p-EGFR, pY-1068, green). Lower panels: staining for ST6GAL1 (red) and total EGFR (t-EGFR, green). Nuclei were stained with Hoechst. Note that the acinar cells of WT mice lack detectable p-EGFR, whereas weak activation of EGFR is noted in ductal cells (arrow). Contrarily, SC acinar cells, as well as nonmalignant KSC acinar cells, display strong EGFR activation. Scale bar = 25µM. (B) Upper panels: PanIN lesions from KC and KSC mice express high levels of ST6GAL1 (red) and p-EGFR (green). Lower panels: PanINs were co-stained for ST6GAL1 (red) and t-EGFR (green). Nuclei were stained with Hoechst. Scale bar = 25µM. (C) Organoids were lysed and immunoblotted for p-EGFR (pY-1068), t-EGFR, and ST6GAL1. (D) Densitometric analyses of ST6GAL1 expression from 3 independent organoid lysates. Densitometric units (DU) for ST6GAL1 were normalized to values for β-tubulin. *p<0.05. (E) Densitometric analyses of p-EGFR, t-EGFR and the p-EGFR/t-EGFR ratio measured from 3 independent organoid lysates. DUs for each target gene were normalized to values for β-tubulin. *p<0.05.

## Discussion

Cancer-associated changes in glycosylation have been reported for decades, and certain glycan structures, such as sialyl Lewis a (sLe^a^, CA19-9), are used in the clinic to monitor cancer progression and recurrence (53). Nonetheless, the potential for glycans and glycosyltransferases to serve as biomarkers or therapeutic targets remains an under-investigated area of cancer research. A role for aberrant glycosylation, including altered sialylation, in pancreatic cancer has been previously documented. Mice engineered to express sLe^a^ in combination with KRAS^G12D^ developed more aggressive pancreatic cancer when compared with mice expressing KRAS^G12D^ alone (54). Sialylation by ST6GAL1 has likewise been implicated in pancreatic cancer. Lectin microarray analyses revealed that α2,6 sialylation was one of the most prominently elevated glycan structures in the neoplastic tissues of PDAC patients and KC mice (55). Deletion of *St6gal1* in KC mice impeded PanIN formation (55). Hsieh *et al*. observed that administration of fructose to cerulein-treated KC mice accelerated invasive PDAC, and ST6GAL1 activity was central to this process (15). The present investigation adds to this body of work by firmly establishing a tumor-promoting function for ST6GAL1 in tumor xenograft models using human PDAC cells, and GEM models with transgenic ST6GAL1 expression in the pancreas. Indeed, the combined expression of ST6GAL1 and KRAS^G12D^ reduced median survival to 4.3 months, an interval comparable to the 5 month survival time of the aggressive “KPC” PDAC model, which harbors dual expression of KRAS^G12D^ and mutant p53 (*Trp53*^*R172H*^) (43).

ST6GAL1 endows cancer cells with stemness features including tumor-initiating potential in limiting dilution assays (17); we therefore postulated ST6GAL1 may facilitate early stages in PDAC development. Results herein show that ST6GAL1 expression is upregulated in Stage I PDAC, with levels increasing in advanced-stage malignancy. The premalignant PanIN lesions of KC mice also display elevated ST6GAL1 expression. Alexander *et al*. similarly observed a progressive upregulation of ST6GAL1 during the transition between gastric premalignancy and gastric cancer, and in this same report, ST6GAL1 was identified as a novel marker for gastric stem cells (29). ST6GAL1 contributes functionally to a number of stemness-associated cellular processes including EMT, iPSC reprogramming, and acquisition of CSC characteristics (17, 23, 27, 28, 31). In the Suit2 and S2-LM7AA PDAC cell lines, high ST6GAL1 activity was shown to induce EMT, and EMT was driven by the ST6GAL1-dependent activation of EGFR in these models (23). RNA-Seq results comparing SC and WT pancreata highlighted a role for ST6GAL1 in activating stemness pathways including Wnt, Notch and Hedgehog. These same pathways were activated when ST6GAL1 was overexpressed in Suit2 PDAC cells (23). The Wnt, Notch and Hedgehog cascades are well-known players in PDAC development (45, 56, 57), and Notch cooperates with oncogenic KRAS in directing ADM and PanIN formation (58). As well, signaling by Wnt and Notch during ADM suppresses the expression of acinar-specific genes (3). The Notch pathway is activated in acinar cells downstream of EGFR signaling. The EGFR-mediated activation of MAPKs in ADM-like cells induces an upregulation in Protein Kinase D1 (PRKD1), which in turn, activates the Notch signaling node (3).

The source of stem/progenitor cells in the adult exocrine pancreas remains controversial, and a *bona fide* stem cell population has yet to be defined. Alternatively, acinar cells are highly plastic, and some acinar subpopulations function as facultative progenitors when tissue regeneration is needed (59, 60). Acinar cells undergoing ADM are a major source of facultative progenitors, and these cells also serve as tumor-initiating cells (59, 60). Here we show that endogenous ST6GAL1 is upregulated in the ADM-like cells induced by cerulein-stimulated pancreatitis, highlighting ST6GAL1 as a new marker for cells undergoing ADM. Moreover, SC mice form quantitatively more ADM-like cells during pancreatitis than WT mice, suggesting a causal role for ST6GAL1 in promoting inflammation-associated ADM. Importantly, the forced expression of ST6GAL1 in acinar cells induces a pronounced upregulation in a ductal gene network (in the absence of either pancreatitis or the KRas oncogene), as evidenced by RNASeq results and staining of SC tissues for SOX9, KRT8 and KRT19. These data align with our prior studies indicating that ST6GAL1 activity enhances SOX9 expression in many cancer cell models (17). The concept that ST6GAL1 reprograms acinar cells to adopt more ductal/progenitor-like characteristics is reinforced by the finding that ST6GAL1 promotes the expression of SOX9 and HES1, while suppressing PTF1A, in both the classic 266-6 ADM cell line, as well as in organoid models. Additionally, ST6GAL1 activity augments the initial formation of organoids as well as organoid growth over time. Clonogenic growth of primary cells in organoid models is a key feature of stem cell behavior (61). Knockdown of ST6GAL1 in KC organoids inhibits organoid formation and growth, and suppresses SOX9 expression while increasing levels of PTF1A. These data underscore the importance of ST6GAL1 in maintaining stem/ductal characteristics in cells with oncogenic KRas. It has long been appreciated that only a subset of KRAS^G12D^-expressing acinar cells within KC mice is susceptible to oncogenic transformation (62, 63). The de-differentiation of acinar cells during the process of ADM renders these cells vulnerable to Kras-driven neoplasia (64, 65).

ST6GAL1 regulates intracellular signaling cascades, and consequently, gene expression, by sialylating select cell surface receptors. ST6GAL1 activity enhances survival-associated pathways including those stimulated by EGFR (24, 66). The activation of EGFR by TGFα is one of the chief mechanisms responsible for inducing ADM (36, 37) and EGFR signaling promotes the expression of SOX9 (35). EGFR signaling is important for the transdifferentiation of acinar cells into ductal-like cells both *in vitro* (47) and *in vivo* (38, 39). EGFR signaling is also essential for KRAS-driven PDAC development (67, 68). EGFR activates wild-type KRAS, and thereby collaborates with mutant *KRAS* alleles to amplify KRAS-dependent pathways (2). Deletion of *Egfr* in the acinar cells of KRAS^G12D^-expressing mice prevents PanIN and PDAC formation (67, 68). We previously reported that ST6GAL1 sialylates and activates EGFR in multiple PDAC and ovarian cancer cell lines (22, 23), although an inhibitory effect of sialylation on EGFR has also been reported (69, 70). More recently, we determined that ST6GAL1-mediated sialylation of EGFR prolongs downstream survival signaling by enhancing EGFR retention on the cell surface (24). It is well-known that *N*-glycans play seminal roles in modulating EGFR structure and signaling (71-73). For example, an *N*-glycan attached to Asn-579 is a central player in maintaining the autoinhibitory tether within the EGFR structure (74), and this *N*-glycan is a known target for sialylation by ST6GAL1 (75). Deletion of this *N*-glycan releases the tether, leading to EGFR activation (74). It is tempting to speculate that the addition of the bulky, negatively-charged sialic acid to this *N*-glycan may interfere with formation of the autoinhibitory tether. While future research will be needed to delineate the specific effects of α2,6 sialylation on the overall structure of EGFR, it is noteworthy that the acinar cells within SC pancreatic tissues, as well as SC-derived organoids, exhibit a dramatic enrichment in EGFR activation when compared to WT acinar cells and organoids. These data point to a potential mechanism by which ST6GAL1-mediated receptor sialylation promotes the upregulation of stem and ductal gene networks.

In summary, results from this investigation suggest that ST6GAL1 activity reprograms mature acinar cells to adopt a more stem and ductal-like phenotype that poises cells for ADM. When occurring in conjunction with an oncogenic event, such as a *KRAS* mutation, this may facilitate neoplastic transformation, as well as the transition to malignancy. An extensive literature has documented ST6GAL1 overexpression in a plethora of human cancers; however, knowledge regarding ST6GAL1’s functional contribution to carcinogenesis remains limited. The collective findings in this report provide new insights into the mechanisms by which ST6GAL1 imparts cell characteristics that foster neoplastic development.

## Methods

### ST6GAL1 IHC staining in human pancreatic tissues

Tissue microarrays containing human nonmalignant and PDAC patient pancreatic tissues were obtained from US Biomax Inc. (PA1001c, PA2072a, PA1921, and PA1002b). Sections were subjected to antigen retrieval using Antigen Unmasking Solution, Citric Acid Based (Vector Laboratories), blocked with 2.5% horse serum for 1 hr, and then incubated overnight at 4°C with goat polyclonal antibody to ST6GAL1 (R&D Systems) (see Table S3 for antibody information). Sections were incubated with ImmPRESS® - HRP anti-goat IgG reagent for 1 hr and developed using ImmPACT® NovaRED or ImmPACT® DAB (Vector Laboratories). Images were captured with a Nikon 80i Eclipse microscope and processed with NIS-Elements Imaging Software.

### Cell cultures

Suit2 and S2-013 cell lines were a kind gift from Dr. Michael A. Hollingsworth at the University of Nebraska. The S2-LM7AA line was developed by Drs. Lacey McNally and Donald Buchsbaum at the University of Alabama at Birmingham. Suit2 cells were transduced with *ST6GAL1*-expressing lentivirus (Genecopoeia, cat#LPP-M0351-Lv105-200-5) or empty vector lentivirus (Sigma). The S2-013 and S2-LM7AA lines were transduced with lentivirus encoding either shRNA against *ST6GAL1* (Sigma, cat#TRCN00000035432) or a control shRNA (Sigma, cat#SHC002V). Murine 266-6 cells (ATCC cat#CRL-2151) were transduced with lentivirus encoding *St6gal1* (Genecopoeia, cat#LPP-EX-Mm05221-Lv105) or shRNA against *St6gal1* (Sigma, cat#TRCN00000018818). Stable polyclonal populations were isolated by puromycin selection.

### Tumor xenograft experiments using the Suit2 isogenic series

Luciferase was stably expressed in Suit2, S2-013, and S2-LM7AA cells using lentivirus (Cellomics Technologies). For the Suit2 and S2-LM7AA lines, a suspension of 1×10^6^ cells in 20 μL of PBS was injected into the pancreas of athymic nude mice (Jackson Laboratories, cat#002019). Tumor growth was monitored by BLI using the IVIS Lumina Series III (Perkin Elmer). At the endpoint (4 weeks), extracted livers were imaged by BLI. For the S2-013 line, 1×10^6^ cells were suspended in 500 μL PBS containing 10% Matrigel, and injected into the flank. Tumor growth was monitored by BLI and tumor volume calculated from caliper measurements using the formula L x W x H. At the endpoint (5 weeks), excised tumors were weighed and lungs imaged by BLI. For the Suit2 and S2-LM7AA experiments, n=7 mice/group, and for the S2-013 line, n=11 mice/group (cohorts included male and female animals). All procedures were performed with prior approval from the UAB Institutional Animal Care and Use Committee (IACUC).

### Development of GEM models

To develop the SC model, an *LSL-ST6GAL1* transgene was expressed under the control of the *Rosa26* gene promoter and inserted in the *Rosa26* locus of C57BL/6 mice as previously described (17). Mice expressing the *LSL-ST6GAL1* transgene were crossed to *Pdx1-Cre* mice. KC mice were obtained by crossing mice expressing *LSL-KRas*^*G12D*^ with the *Pdx1-Cre* line. KSC mice were obtained by crossing SC (*Rosa26-LSL-ST6GAL1; Pdx1-Cre*) mice with *LSL-KRas*^*G12D*^ mice. Genotype was verified by PCR using the following primers:

Kras mutant forward: 5’ – AAGCTAGCCACCATGGCT - 3’

Kras Reverse: 5’ – CGCAGACTGTAGAGCAGCG - 3’

Cre Forward: 5’ - TGCCACGACCAAGTGACAGC - 3’

Cre Reverse: 5’ - CCAGGTTACGGATATAGTTCATG - 3’

ST6GAL1 Forward: 5’ – CCAGGACCAGGCATCAAGTT – 3’

ST6GAL1 Reverse: 5’ – CCCATAGCTCCCAAGGCATC – 3’

ST6GAL1 expression in SC mice was confirmed by IHC, and increased α2,6 sialylation was validated by staining cells with SNA. Briefly, pancreata were digested using Miltenyi tissue dissociation kit (cat#130-110-201), and dispersed cells passed through a 100 μm filter. Cells were resuspended in PBS containing 10µM ROCK inhibitor (Hello Bio cat#HB2297) to improve viability. Cells were stained for 30 min on ice with: Aqua live/dead stain (Life Technologies); anti-EpCAM-PE (BioLegend); and SNA-FITC (Vector Labs). Mean Fluorescent Intensity values for SNA staining were measured on viable cells (Aqua negative) that were positive for EpCAM (epithelial marker). Unstained and single-color controls were used for compensation and gating, and only viable singlets were included. An LSRII was used to acquire data and FlowJo V10 was used for data analysis. Experiments were conducted on 3 mice per group.

For survival analyses (n=10 mice/group), KSC and KC mice were euthanized per IACUC guidelines when they lost 20% of their body weight and/or presented with low body score. Euthanized mice were evaluated for macro-metastases in lungs, liver, intestines, peritoneum and other organs, and micro-metastases were identified by examining H&E-stained tissue sections.

### Generation and culture of organoid lines

As previously described (49), pancreatic tissues were minced and placed in collagenase I solution for 45 min with intermittent trituration. The tissues were then passed through a 70 μm cell strainer. Cells were washed with media and resuspended in Matrigel (Corning, cat#CB-40234). 15 μL of the cell suspension was placed in a 24-well plate. The plate was then inverted and the Matrigel allowed to solidify at 37°C for 12 min. After solidification, cultures were placed in organoid media and allowed to grow for 3 days. Organoid cultures were passaged until they were free of contaminating cells such as fibroblasts, and then used for experiments. As described (76), organoids were propagated in DMEM containing 50% L-WRN-conditioned media with the following supplements: 10 µM ROCK inhibitor (Hello Bio cat#HB2297); 10 µM TGFβR1 inhibitor (Selleck Chemicals cat#S1067); and 10 mM nicotinamide (LKT Labs cat#N3310).

For some experiments, organoid-derived cells were placed in adherent monolayer culture. Wells were coated with Matrigel (1:30 dilution in PBS), and the Matrigel was allowed to solidify for 30 min. Organoids were dissociated and seeded onto the Matrigel-coated wells in media composed of DMEM, 10% FBS, 5% organoid media and pen/strep supplement. Monolayer cultures were grown for 72 hr and lysed for RNA extraction. To evaluate organoid-forming potential, 2000 organoid-derived cells were plated in 10µl of Matrigel and allowed to solidify. The cultures were grown in organoid media for 3 days, and the total number of organoids within each well was counted. To monitor organoid growth over time, 2000 cells were seeded into organoid culture and at days 3, 6 and 9, organoid cultures were harvested and dissociated into single cell suspensions using trypsin. Live cells (that exclude Trypan blue) were counted using a hemocytometer. At least 3 biological replicates (i.e., 3 distinct organoid cultures) were analyzed.

KC organoids with ST6GAL1 knockdown (KC-KD) were generated by transducing dissociated organoid cells with lentivirus encoding shRNA against *St6gal1*, using the vector described above for 266-6 cells. Control organoids (KC-shC) were generated using lentivirus encoding a nontargeting shRNA (cat#SHC005V). ∼50,000 organoid cells were mixed with the lentivirus at an MOI of 5, and incubated at 37°C for 6 hr. After washing with media to remove lentiviral particles, cells were resuspended in Matrigel and plated to allow re-formation of the organoids. Stably transduced organoids were selected with puromycin for two days post-transduction.

### RNA extraction and qRT-PCR

RNA was extracted using the Ambion RNA extraction kit (Life Technologies). The M-MLV reverse transcriptase (Promega) was used for cDNA preparation. qRT-PCR was conducted using the Taq-Man fast master mix (Thermo) using primers for *Sox9* (Mm00448840_m1), *Hes1* (Mm01342805_m1), *Ptf1a* (Mm00479622_m1), *Lgr5* (Mm000438890_m1) and *Axin2* (Mm00443610_m1). *Gapdh* primers (Mm99999915_g1) were used for normalization. At least 3 independent experiments were performed, with each qRT-PCR experiment performed in triplicate.

### Immunoblotting

Cell lines were lysed in RIPA buffer containing protease and phosphatase inhibitors (Thermo Fisher). Organoids were centrifuged at 100xg, incubated in Accutase for 15 min at 37°C, and then the dissociated cells were lysed in RIPA buffer with protease and phosphatase inhibitors. Protein concentration was assessed by BCA (Thermo Fisher). Following SDS-PAGE, proteins were transferred to PVDF membrane. Membranes were blocked in 5% nonfat dry milk, and incubated with primary antibody overnight at 4°C. Primary antibodies included anti-SOX9 (AbCam), anti-HES1 (AbCam), anti-PTF1A (AbCam), anti-EGFR (Cell Signaling Technology), anti-phospho-EGFR (pTyr-1068, Cell Signaling Technology), and anti-ST6GAL1 (R&D systems). Membranes were incubated in secondary antibody for 1 hr, and developed using Clarity Western ECL substrate (BioRad, cat# 1705061). Densitometry was conducted using ImageJ.

### Cerulein treatment and flow cytometric analysis of ADM cells

Pancreatitis was induced by 7 intraperitoneal injections of cerulein (50 μg/kg; Tocris Bioscience, cat#62-641) at hourly intervals on two alternating days. Saline injections were used as the control. On day 4, pancreata were isolated and digested for 30 min using the Miltenyi tissue dissociation kit (cat#130-110-201) supplemented with a ROCK inhibitor. Digested tissue was passed through a 100 μm filter and centrifuged at 300xg for 5 min. The pellet was resuspended in PBS with ROCK inhibitor. Under these conditions, the viability of dissociated cells was >90%. The cell suspension was stained with Aqua live/dead stain (Life Technologies), anti-EpCAM-PE (BioLegend), anti-CD45-APC (BioLegend), anti-CD133-PE/Cy-7 (BioLegend) and UEA-FITC (Sigma-Aldrich) for 30 min. UEA and anti-CD133 staining was quantified on cells positive for EpCAM (epithelial marker), negative for CD45 (immune marker), and negative for the Aqua dye. Unstained and single color controls were used to set voltage and compensation, and only viable singlets were counted. Experiments were conducted on 3 mice per group.

### Histological Analyses

#### IHC

Tissues from GEM models were processed for antigen retrieval and IHC staining as described above. Primary antibodies included anti-ST6GAL1 (R&D systems) and anti-SOX9 (AbCam). Slides were counterstained with haematoxylin (Vector Labs), dehydrated and mounted with coverslips using VectaMount medium (Vector Labs).

#### Immunofluorescence staining

Paraffin-embedded tissues were processed as above and incubated in primary antibody overnight 4°C. Primary antibodies included anti-SOX9 (AbCam), anti-EGFR (AbCam), anti-phospho-EGFR (pTyr-1068, AbCam), anti-GM130 (AbCam), anti-pancreatic α Amylase (AbCam), anti-KRT8 (AbCam), anti-KRT19 (AbCam), and anti-ST6GAL1 (R&D systems). Slides were incubated with the appropriate secondary antibody, followed by incubation in Hoechst solution in PBS for 5 min at RT. Slides were mounted using VECTASHIELD® Vibrance™ Antifade Mounting Medium (Vector Labs).

#### H&E, Alcian Blue and Sirius Red staining

For H&E staining, slides were exposed to haematoxylin for 7 min (VitroVivo), then immersed in 95% alcohol for 1 min, followed by eosin staining for 30 sec. H&E stained slides from 20 week-old KC and KSC mice were analyzed for PanINs and adenocarcinoma under blinded conditions by a board-certified pathologist, Dr. Isam-Eldin Eltoum. Tissues were also stained with: (i) Alcian Blue (IHC World) followed by counterstaining with Nuclear Fast Red (Vector Labs), and (ii) Sirius Red (IHC world) followed by counterstaining with haematoxylin. Staining was quantified using a stereological approach. A grid was placed over an image of the entire pancreas, and the intersection points landing on tissues positively stained for either Alcian Blue or Sirius Red were counted. To obtain the stereologic score, the number of intersection points landing on positively-stained tissues was divided by the total number of intersection points counted for the slide.

### RNA Sequencing

Pancreata from 20 week-old mice (3 mice/genotype) were extracted and placed in RNAlater (Invitrogen) overnight at 4°C. Samples were then transferred into TRIZOL (Thermo, cat#15596026) and homogenized. 100 µL chloroform was added for each mL of TRIZOL. The top aqueous layer was isolated, and an equal volume of 70% ethanol added. The protocol from the Qiagen RNeasy Mini Kit was subsequently followed to isolate the RNA (Qiagen, cat# 74104). RNA sequencing was conducted using the Illumina NextSeq500 (Illumina Inc) at the UAB Hugh Kaul Genomics Core. RNA quality was assessed using the Agilent 2100 Bioanalyzer, and RNA with an RNA Integrity Number (RIN) of ≥7.0 was used for library preparation. RNA was converted to a sequencing ready library using the NEBNext Ultra II Directional RNA library kit (New England Biolabs). The cDNA libraries were quantitated using qPCR in a Roche LightCycler 480 with the Kapa Biosystems kit for Illumina library quantitation (Kapa Biosystems). Cluster generation was performed according to the manufacturer’s recommendations for onboard clustering (Illumina). We generated 30-35 million paired-end 75bp sequencing reads per sample for gene level abundance.

#### Quantification of gene expression and differential expression

Pre-alignment quality assessments of the raw fastq sequences were carried out using FastQC (version 0.11.7, http://www.bioinformatics.babraham.ac.uk/projects/fastqc). STAR (version 2.7.3a) was used to align the raw RNA-seq fastq reads to the mouse reference genome (GRCm38 p4 Release M11) from Gencode (77). Following alignment, HTSeq-count (version 0.11.3) was used to count the number of reads mapping to each gene (78). Normalization and differential expression were then applied to the count files using DESeq2 (version 1.26.0) using default parameters in their vignette (79).

#### Ingenuity Pathway Analysis (IPA)

A data set containing gene identifiers and corresponding expression values was uploaded into IPA (https://digitalinsights.qiagen.com/products-overview/discovery-insights-portfolio/analysis-and-visualization/qiagen-ipa/). Each identifier was mapped to its corresponding object in Ingenuity’s Knowledge Base (80). A fold change cutoff of ±2 and p value < 0.05 was set to define Network Eligible molecules. These molecules were overlaid onto a global molecular network developed from information contained in Ingenuity’s Knowledge Base. Networks of Network Eligible Molecules were then algorithmically generated based on their connectivity.

#### Gene set enrichment analysis (GSEA)

GSEA was performed on the full normalized gene expression data using GSEA software version 4.1.0 and the Molecular Signature Database (MSigDB) (version 7. 2) (81, 82). Default parameters were used except that the permute parameter was set to the gene set and plot graphs for the top sets of each phenotype set to 50.

### Statistical Analyses

Quantification of ST6GAL1-positive cells in human pancreatic specimens was conducted using the nonparametric Kruskal-Wallis test followed by Dunn’s multiple comparison test. A nonparametric test was selected after evaluating the data with the D’Agostino-Pearson normality test. For the tumor xenograft studies, BLI experiments were analyzed using a Two-Way ANOVA, and tumor volume measurements were evaluated using a Mann-Whitney test. Mean survival time for KC and KSC mice was calculated using a Kaplan-Meier statistical analysis. For other experiments, a two-tailed Student’s t-test was used to evaluate the data, where p < 0.05 was considered as significant. Graphs depict mean ± S.E.M.

## Supplementary Material

Representative images of ST6GAL1 expression in human PDAC patient tissues are provided in Fig. S1. Fig. S2 includes SNA staining of the Suit2 isogenic cell series as well as histological analyses conducted on Suit2, S2-LM7AA and S2-013-derived tumors. Fig. S3 includes H&E and ST6GAL1 IHC staining of tissues from GEM models. Table S1 provides extended data from GSEA analyses comparing SC and WT mice. Table S2 provides extended data from GSEA analyses comparing KSC and KC mice. Table S3 lists the antibodies and lectins used in this study.

## Data Availability

All of the data are contained within the manuscript or supplementary figures, with the exception of the RNA-Seq results. The RNA-Seq data have been deposited in NCBI’s Gene Expression Omnibus (GEO) and are accessible through GEO Series accession number GSE169546.

## Author Contributions

AC, NB, MPM, JH, CMB, RBJ, DKC, and MRC contributed to data curation. AC, NB, MPM, CMB, DKC, IEE and SLB were involved in formal analysis of the data. CAK and DJB provided resources, and AC, NB, MPM, CMB, KLA, LES, PDS, BS, LEH, MMB, CSH and ZY assisted in developing the methodology. Composition of the original draft was executed by AC, NB, and SLB, and the manuscript was edited and reviewed by all authors. SLB was responsible for project administration and funding acquisition.

## Acknowledgements

This study was supported by National Institutes of Health grant U01 CA233581, Alliance of Glycobiologists for Cancer Research (SLB), and a grant from the Richard A. Elkus Foundation for GI Oncology Research (SLB). MPM was supported by National Institutes of Health grant F32 CA264906. The authors gratefully acknowledge assistance from the following Shared Facilities at UAB: Small Animal Imaging Shared Facility; Transgenic Mouse Core; Flow Cytometry Core; Heflin Genomics Core Laboratory, and the Stem Cell Organogenesis Program.

## Notes

The authors declare no potential conflicts of interest

### Competing Interest Statement

The authors have declared no competing interest.

