## Supplementary Figures and Tables for "ST6GAL1 sialyltransferase promotes acinar to ductal metaplasia and pancreatic cancer progression"

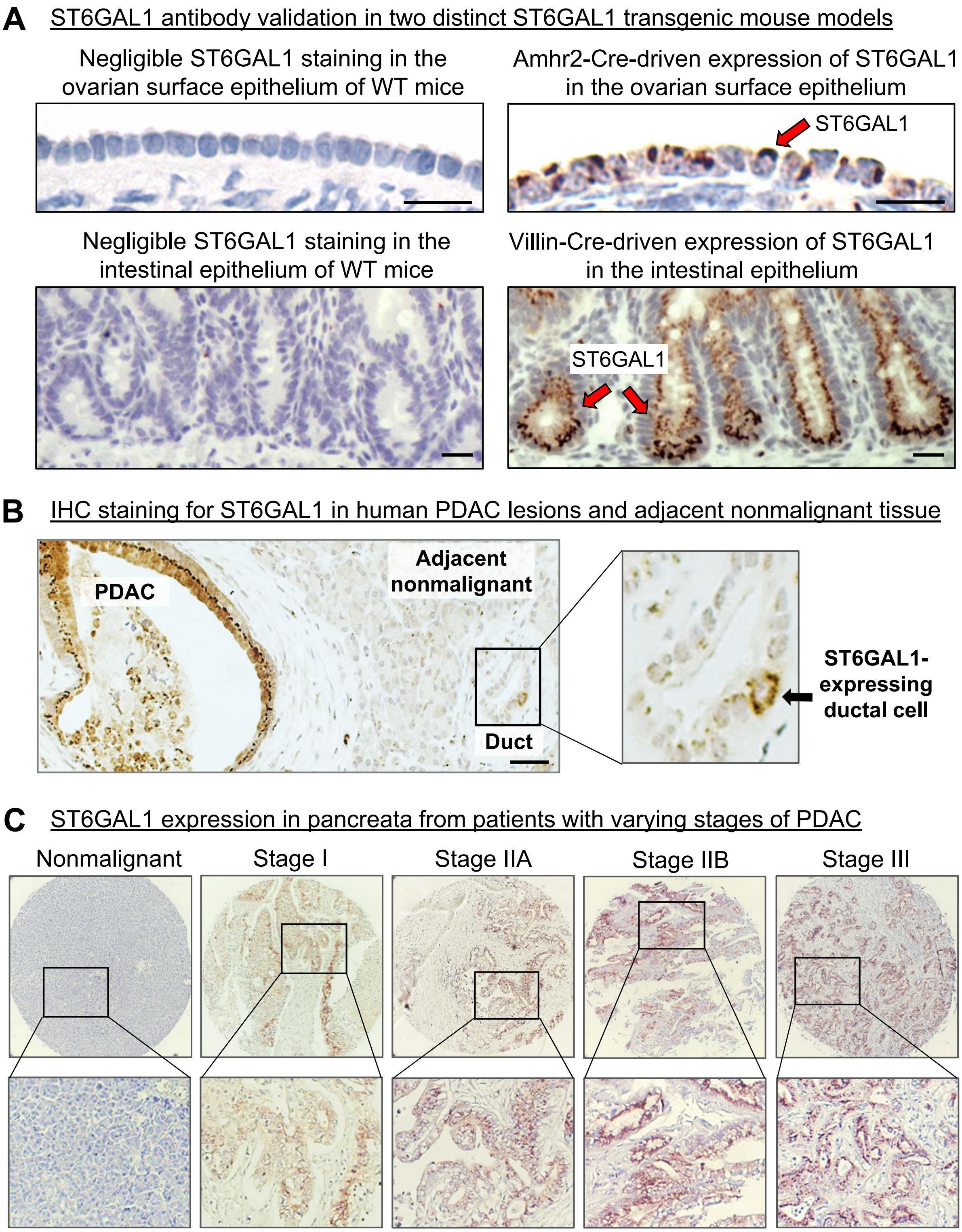


**Fig. S1. ST6GAL1 expression in nonmalignant and malignant patient pancreatic tissues**

(A) The ST6GAL1 antibody was validated by IHC staining in two GEM models with ectopic expression of ST6GAL1 (note that this antibody recognizes both human and murine ST6GAL1). Strong ST6GAL1 staining was noted in the ovarian epithelium of mice with Amhr2-Cre driven ST6GAL1 expression (top right panel), as well as in the intestinal epithelium of mice with Villin-Cre driven ST6GAL1 expression (lower right panel). Scale bar = 20 μM.

(B) IHC staining for ST6GAL1 in a PDAC patient sample. ST6GAL1 is extensively expressed throughout the PDAC lesion, whereas expression is negligible in the nonmalignant, adjacent pancreatic tissue. The inset depicts a rare, ST6GAL1-positive ductal cell within the nonmalignant adjacent tissue. Scale bar = 50 μM.

(C) ST6GAL1 IHC staining in representative specimens from patients with varying stages of PDAC.

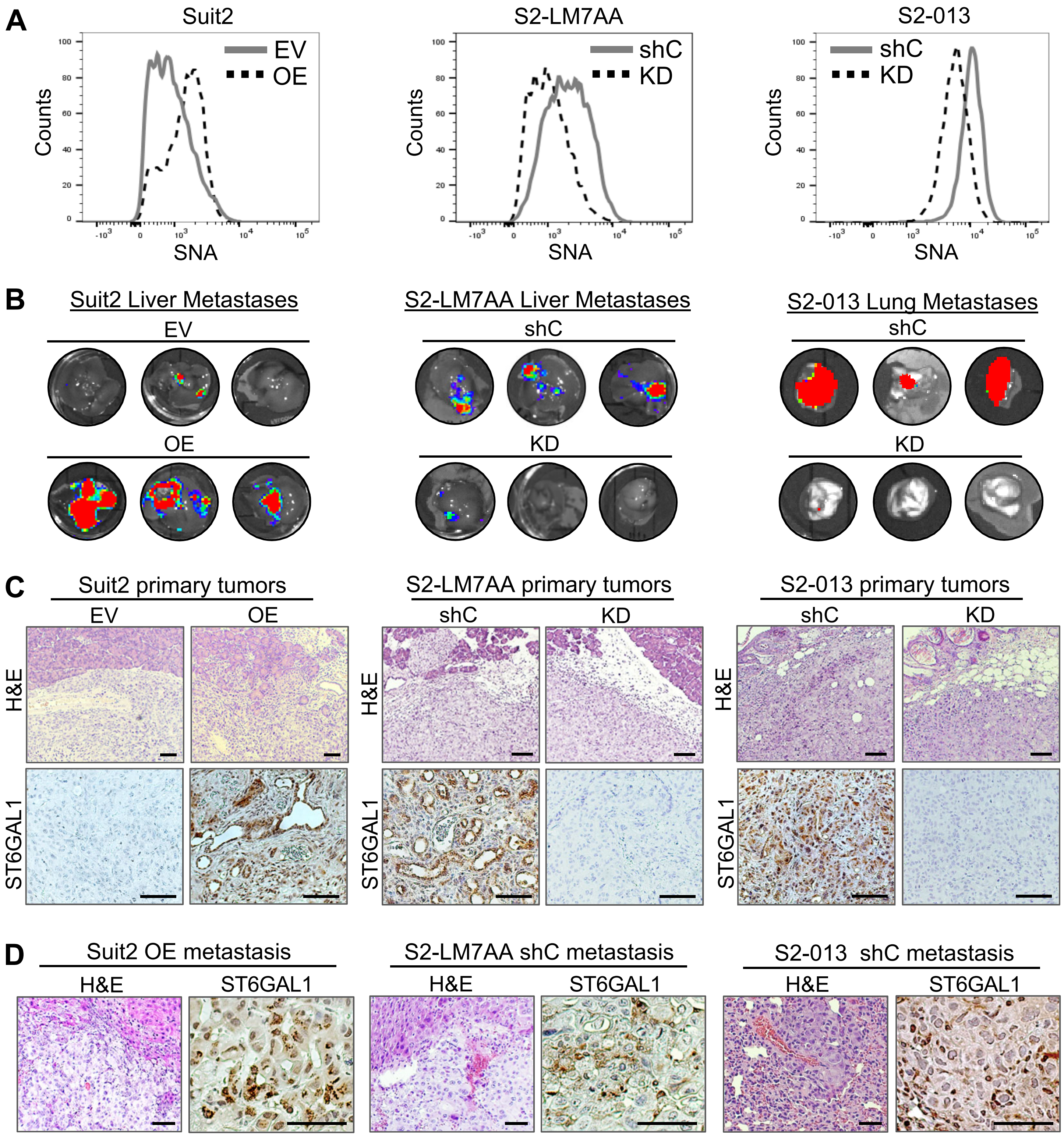


**Fig. S2. Tumor xenografts using the isogenic human Suit2 PDAC cell series.**

(A) Cells with modulated ST6GAL1 expression (overexpression, OE, or knockdown, KD) were stained with the SNA lectin, which binds α2,6 sialic acid, and then analyzed for surface sialylation by flow cytometry.

(B) Bioluminescence imaging (BLI) of representative organs harboring metastatic tumors formed from Suit2, S2-LM7AA and S2-013 cells. Each image is from a distinct mouse.

(C) Upper panels: H&E stained primary tumors from the Suit2, S2-LM7AA and S2-013 cohorts. Lower panels: IHC staining for ST6GAL1 on primary tumors from the Suit2, S2-LM7AA and S2-013 cohorts. Scale bar = 100 μM

(D) H&E (left panels) or IHC staining for ST6GAL1 (right panels) on metastatic tumors from the cell lines with high ST6GAL1 expression (Suit2 OE; S2-LM7AA shC; S2-013 shC). Scale bar = 50 μM

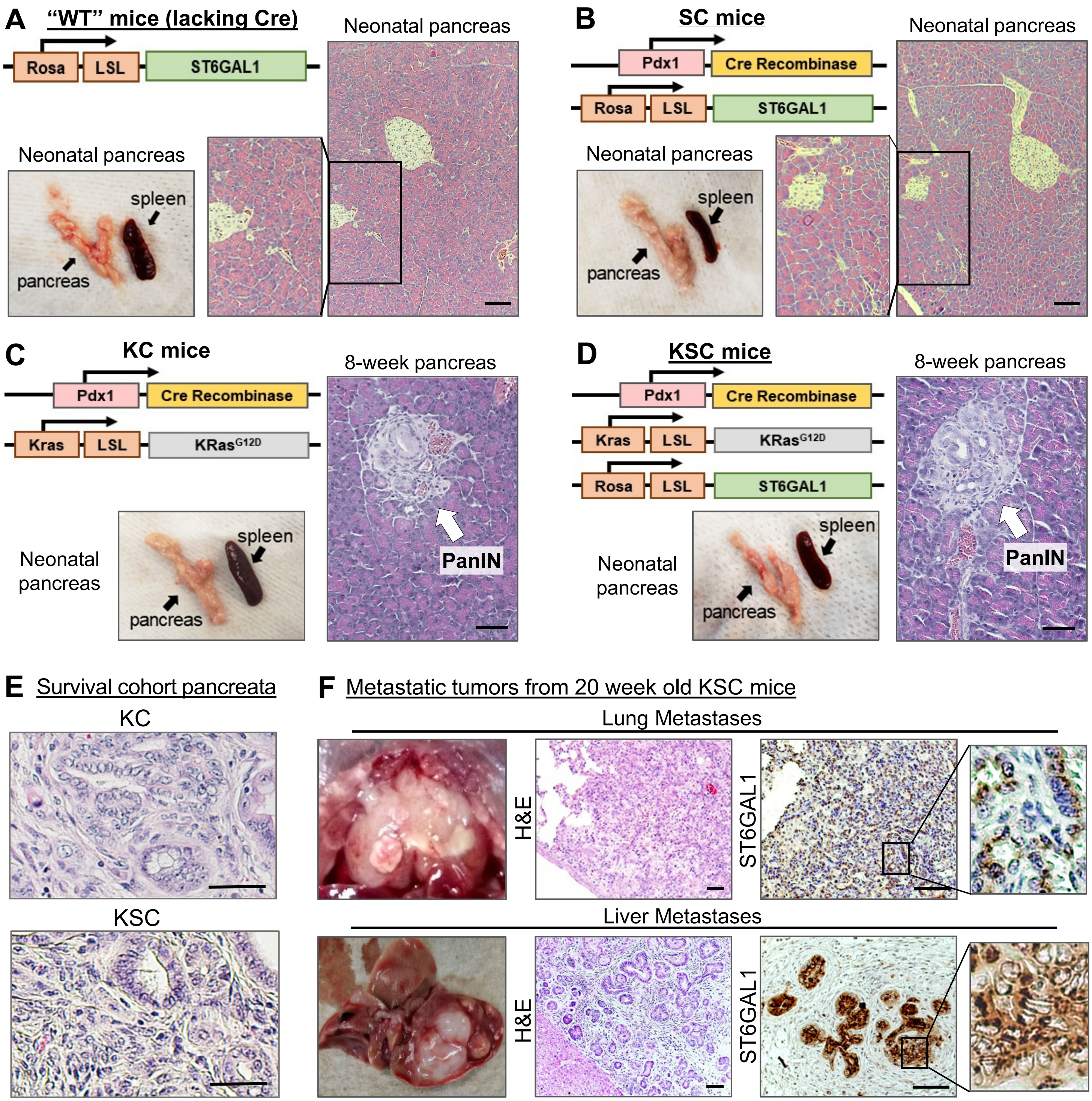


**Fig. S3. Histologic evaluation of tissues from GEM models**

(A) Image of whole pancreas and H&E stained pancreatic tissues from neonatal mice expressing the *ST6GAL1* transgene, but not the Cre recombinase, abbreviated as "WT" mice. Scale bar = 100 μM.

(B) Image of pancreas and H&E stained tissues from neonatal SC mice (expressing the *ST6GAL1* transgene plus Cre recombinase). No abnormalities were detected in neonatal SC pancreata. Scale bar = 100 μM.

(C) Image of pancreas and H&E stained tissues from KC mice. No pancreatic abnormalities were noted in neonatal mice, however PanIN lesions were detected in 8-week old mice. Scale bar = 50 μM

(D) Image of pancreas and H&E stained tissues from KSC mice. No pancreatic abnormalities were observed in neonatal mice, however PanIN lesions were apparent in 8-week old mice. Scale bar = 50 μM.

(E) H&E stained pancreata from KC and KSC mice in the survival cohort. Scale bar = 50 μM.

(F) Metastatic tumors from 20 week old KSC mice. Upper panels depict lung metastases. Lower panels depict liver metastases: Scale bar = 100 μM.

**Table S1. GSEA analyses of pathways upregulated in SC mice vs. WT mice**

| PATHWAYS | NES | FDR |
|---|---|---|
| ***Developmental/Stemness Pathways*** | | |
| GO EXOCRINE SYSTEM DEVELOPMENT | 2.33 | 0.001 |
| GO CELL MORPHOGENESIS | 2.21 | 0.002 |
| HALLMARK WNT BETA CATENIN SIGNALING | 1.59 | 0.018 |
| GO BETA CATENIN TCF COMPLEX ASSEMBLY | 2.10 | 0.006 |
| PID HES HEY PATHWAY | 1.57 | 0.089 |
| REACTOME REGULATION OF RUNX1 EXPRESSION AND ACTIVITY | 2.01 | 0.015 |
| PID BMP PATHWAY | 1.82 | 0.021 |
| WP HIPPOYAP SIGNALING PATHWAY | 1.59 | 0.086 |
| ***Growth Factor Receptor Signaling Pathways*** | | |
| GO TRANSMEMBRANE PROTEIN RECEPTOR KINASE ACTIVITY | 2.15 | 0.004 |
| REACTOME SIGNALING BY RECEPTOR TYROSINE KINASES | 1.74 | 0.050 |
| WP EGFR TYROSINE KINASE INHIBITOR RESISTANCE | 1.83 | 0.027 |
| REACTOME SIGNALING BY ERBB2 IN CANCER | 1.88 | 0.027 |
| REACTOME MET PROMOTES CELL MOTILITY | 2.08 | 0.009 |
| REACTOME SIGNALING BY FGFR1 IN DISEASE | 2.21 | 0.004 |
| PID VEGFR1_2 PATHWAY | 1.83 | 0.021 |
| REACTOME SIGNALING BY PDGF | 2.16 | 0.004 |
| ***Signaling by Small G Proteins*** | | |
| GO SMALL GTPASE BINDING | 2.43 | 0.001 |
| GO RAS GUANYL NUCLEOTIDE EXCHANGE FACTOR ACTIVITY | 2.24 | 0.002 |
| PID RHOA REG PATHWAY | 2.00 | 0.009 |
| GO RAB GTPASE BINDING | 2.44 | 0.001 |
| ***Cell Adhesion and Integrin Signaling*** | | |
| PID N-CADHERIN PATHWAY | 1.96 | 0.016 |
| PID INTEGRIN1 PATHWAY | 1.88 | 0.020 |
| PID AVB3 INTEGRIN PATHWAY | 1.84 | 0.020 |
| PID FAK PATHWAY | 1.92 | 0.019 |
| KEGG FOCAL ADHESION | 2.06 | 0.006 |
| ***Cancer Pathways*** | | |
| KEGG PATHWAYS IN CANCER | 1.78 | 0.020 |
| WP BREAST CANCER | 1.93 | 0.017 |
| WP BLADDER CANCER | 1.74 | 0.048 |
| KEGG RENAL CELL CARCINOMA | 1.68 | 0.030 |
| KEGG PROSTATE CANCER | 1.63 | 0.045 |
| KEGG THYROID CANCER | 1.82 | 0.015 |
| KEGG NON SMALL CELL LUNG CANCER | 1.76 | 0.020 |
| KEGG ENDOMETRIAL CANCER | 1.73 | 0.026 |
| KEGG GLIOMA | 1.76 | 0.020 |
| KEGG CHRONIC MYELOID LEUKEMIA | 1.69 | 0.030 |

**Table S2. GSEA analyses of pathways upregulated in KSC mice vs. KC mice**

| PATHWAYS | NES | FDR |
|---|---|---|
| ***Developmental/Stemness Pathways*** | | |
| GO REGULATION OF EMBRYONIC DEVELOPMENT | 2.090 | 0.014 |
| REACTOME TRANSCRIPTIONAL REGULATION OF PLURIPOTENT STEM CELLS | 1.875 | 0.047 |
| GO POSITIVE REGULATION OF STEM CELL PROLIFERATION | 2.223 | 0.011 |
| GO CELL MORPHOGENESIS | 2.036 | 0.014 |
| REACTOME REGULATION OF RUNX1 EXPRESSION AND ACTIVITY | 2.060 | 0.011 |
| PID KIT PATHWAY | 1.649 | 0.045 |
| PID BMP PATHWAY | 1.853 | 0.024 |
| PID HES HEY PATHWAY | 1.616 | 0.052 |
| GO HIPPO SIGNALING | 2.101 | 0.016 |
| ***Growth Factor Receptor Signaling Pathways*** | | |
| GO TRANSMEMBRANE RECEPTOR PROTEIN TYROSINE KINASE ACTIVITY | 1.852 | 0.031 |
| BIOCARTA EGF PATHWAY | 1.939 | 0.024 |
| GO EPIDERMAL GROWTH FACTOR RECEPTOR BINDING | 1.710 | 0.060 |
| BIOCARTA HER2 PATHWAY | 2.029 | 0.045 |
| BIOCARTA MET PATHWAY | 1.656 | 0.076 |
| REACTOME MET PROMOTES CELL MOTILITY | 1.942 | 0.026 |
| WP PDGF PATHWAY | 1.697 | 0.048 |
| WP PDGFRBETA PATHWAY | 1.959 | 0.016 |
| KEGG TGFBETA SIGNALLING PATHWAY | 1.647 | 0.082 |
| ***Signaling by Small G Proteins*** | | |
| HALLMARK KRAS SIGNALING UP | 1.877 | 0.002 |
| PID RHOA REG PATHWAY | 1.665 | 0.042 |
| BIOCARTA RAC1 PATHWAY | 1.660 | 0.079 |
| ***Cell Adhesion and Integrin Signaling*** | | |
| PID N-CADHERIN PATHWAY | 1.715 | 0.034 |
| PID INTEGRIN1 PATHWAY | 1.805 | 0.033 |
| PID AVB3 INTEGRIN PATHWAY | 1.799 | 0.028 |
| GO FOCAL ADHESION ASSEMBLY | 2.074 | 0.010 |
| PID FAK PATHWAY | 1.862 | 0.024 |
| GO FILOPODIUM | 1.595 | 0.079 |
| GO LAMELLAPODIUM | 1.901 | 0.043 |
| GO INVADOPODIUM | 1.768 | 0.035 |
| ***Cancer Pathways*** | | |
| WP GASTRIC CANCER NETWORK | 1.755 | 0.038 |
| WP HEAD AND NECK SQUAMOUS CELL CARCINOMA | 1.801 | 0.031 |
| KEGG SMALL CELL LUNG CANCER | 1.670 | 0.083 |
| KEGG BASAL CELL CARCINOMA | 1.582 | 0.097 |

**Table S3: Antibody and Lectin information**

| Antibody or Lectin | Application | Dilution | Company (Cat#) |
|---|---|---|---|
| ST6GAL1 | IHC, IF | 1:75 | R&D systems (AF5924) |
| Sox9 | IHC, IF | 1:250 | Abcam (ab185230) |
| EGFR | IF | 1:250 | Abcam (ab52894) |
| p-EGFR (Tyr1068) | IF | 1:250 | Abcam (ab40815) |
| GM130 | IF | 1:200 | Abcam (ab52649) |
| Pancreatic alpha amylase | IF | 1:2000 | Abcam (ab199132) |
| KRT8 | IF | 1:100 | Thermo Fisher (PA5-29607) |
| KRT19 | IF | 1:100 | Proteintech (107-12-1AP) |
| EpCAM-PE | Flow cytometry | 1:200 | BioLegend (118205) |
| SNA-FITC | Flow cytometry | 1:200 | Vector laboratories (FL-1301-2) |
| CD45-APC | Flow cytometry | 1:200 | BioLegend (103111) |
| CD133-PE/Cy7 | Flow cytometry | 1:200 | BioLegend (141209) |
| UEA-FITC | Flow cytometry | 1:200 | Sigma (L9006) |
| ST6GAL1 | WB | 1: 500 | R&D systems (AF5924) |
| Sox9 | WB | 1:1000 | Abcam (ab185230) |
| Hes1 | WB | 1:1000 | Abcam (ab71559) |
| Ptf1a | WB | 1:2000 | Abcam (ab182398) |
| EGFR | WB | 1:1000 | CST (4267) |
| p-EGFR (Tyr1068) | WB | 1:1000 | CST (3777) |
